# Large-scale all-optical dissection of motor cortex connectivity reveals a segregated functional organization of mouse forelimb representations

**DOI:** 10.1101/2021.07.15.452461

**Authors:** Francesco Resta, Elena Montagni, Giuseppe de Vito, Alessandro Scaglione, Anna Letizia Allegra Mascaro, Francesco Saverio Pavone

## Abstract

In rodent motor cortex, the rostral forelimb area (RFA) and the caudal forelimb area (CFA) are major actors in orchestrating the control of forelimb complex movements. However, their intrinsic connections and reciprocal functional organization are still unclear, limiting our understanding of how the brain coordinates and executes voluntary movements. Here we causally probed cortical connectivity and activation patterns triggered by transcranial optogenetic stimulation of ethologically relevant complex movements exploiting a novel large-scale all-optical method in awake mice. Results show specific activation features for each movement class, providing evidence for a segregated functional organization of CFA and RFA. Importantly, we identified a second discrete lateral grasping representation area, namely lateral forelimb area (LFA), with unique connectivity and activation patterns. Therefore, we propose the LFA as a distinct motor representation in the forelimb somatotopic motor map.

## INTRODUCTION

Complex motor behaviors are a combination of discrete and rhythmic movements^1–3^. Discrete movements, such as reaching and grasping, are described as continuous movements from a starting to an ending point in space, while rhythmic movements are periodic, repetitive, and stereotyped, like locomotion or licking^1,4^. In rodents, forelimb movements are controlled by two distinct cortical functional areas: the caudal forelimb area (CFA) and the rostral forelimb area (RFA). Although a large number of works elucidated the anatomical structure and functional outputs of these subregions, their intrinsic connections and reciprocal functional role are unclear^5^.

The functional organization of the motor cortex is canonically investigated combining cortical electrical stimulation and behavioral readout to map cortical movement representations^6–11^. However, the electrophysiological approach is invasive, limited in spatial resolution and lacks cellular population selectivity, thus it is being progressively replaced by optogenetic, which overcomes these limitations^4,12–14^. Light-based motor maps (LBMMs) are powerful tools to study motor cortex topography, however important questions remain concerning the mechanisms that coordinate the activity of different functional areas during voluntary movement execution. In particular, the relative contribution and hierarchic relationship of the RFA and CFA in control forelimb movements are still debated. To answer these questions, it would be beneficial to combine motor mapping with large-scale monitoring of cortical activity. Wide-field fluorescence imaging of voltage-sensitive dyes (VSDs) or genetically-encoded calcium indicators (GECIs) represents the most effective strategy to study motor cortex activity at mesoscale level^15–18^. Although VSDs present high temporal resolution (sub-milliseconds), the use of dyes rules out the possibility to target specific cell populations and to perform longitudinal studies, due to the invasive nature of the dye application. Despite the slower kinetics, GECIs overcome these limitations and show a higher signal-to-noise ratio, resulting in the best choice to study cortical connectivity at the mesoscale level in awake mice. Nevertheless, a crucial factor to consider when combining fluorescence imaging and optogenetics in all-optical configurations is the excitation/absorption spectral overlap that leads to crosstalk between imaging and photostimulation^19^.

Here we first established a crosstalk-free large-scale all-optical experimental configuration combining wide-field fluorescence imaging of the red-shifted GECI jRCaMP1a and optogenetic stimulation of Channelrhodopsin-2 (ChR2) to perform light-based motor mapping of complex movements in awake mice while monitoring the cortical activity of excitatory neurons. Thanks to this low-invasive transcranial method, we investigated RFA and CFA effective connectivity and functional dependencies by causally dissecting the cortical activity patterns triggered by optogenetic stimulation. Our results show a modular organization of the movement-specific activated areas and peculiar activity propagation hallmarks during forelimb complex movement execution. Importantly, we identified a discrete lateral and caudal grasping cortical representation expressing distinct topographic and connectivity features.

## RESULTS

To explore the intrinsic connections and reciprocal functional role of RFA and CFA, we implemented a novel large-scale all-optical method to causally dissect the cortical activity patterns triggered by optogenetic stimulation of two ethologically relevant forelimb movements. Initially, we mapped the cortical representation of two distinct complex movements, a grasping-like movement and a locomotion-like movement optogenetically evoked in RFA and CFA respectively. Next, we recorded the mesoscale cortical response during movement execution. Finally, we quantified the activation maps related to the movements somatotopy to study the RFA and CFA connectivity.

### Wide and long-term stable cortical transfection of both jRCaMP1a and ChR2 in mouse motor cortex

In mice expressing optogenetic actuators in the motor cortex, several classes of limb movements can be evoked depending on the stimulated site^4,12,13^. To investigate large-scale cortical activity underlying the optogenetically-evoked movements, we infected the frontal right cortical hemisphere of C57/B6 mice with adeno-associated viruses (AAV) carrying the red-shifted GECI jRCaMP1a and the optogenetic actuator ChR2. The optical setup consisted of a double-path illumination system integrated into a custom-made wide-field fluorescence microscope for parallel laser stimulation and wide-field cortical imaging (fig. 1a). To evaluate the transfection extension and its long-term stability, the spatial fluorescence intensity profiles were calculated (fig. 1b). In line with our previous observations^20^, we found that ChR2 and jRCaMP1a expression covered all the motor cortices, and their expression levels were highly stable over several weeks (fig. 1b; supplementary fig. 3). Moreover, the expression profiles on histological brain slices showed that the transfection of jRCaMP1a and ChR2 were restricted to the motor cortex reaching all the cortical layers (fig. 1d). Besides, to evaluate the synapsin-targeted jRCaMP1a transfection efficiency, brain slices were stained with the neuronal marker NeuN, revealing that the jRCaMP1a^+^ neurons were 70.2 ± 4.9 % of the NeuN^+^ cells (fig. 1e). Therefore, our injection method leads to a wide transfection that covers all the motor cortices and is sufficiently stable to perform experiments several weeks after injection with stable expression levels.

**Fig. 1.**
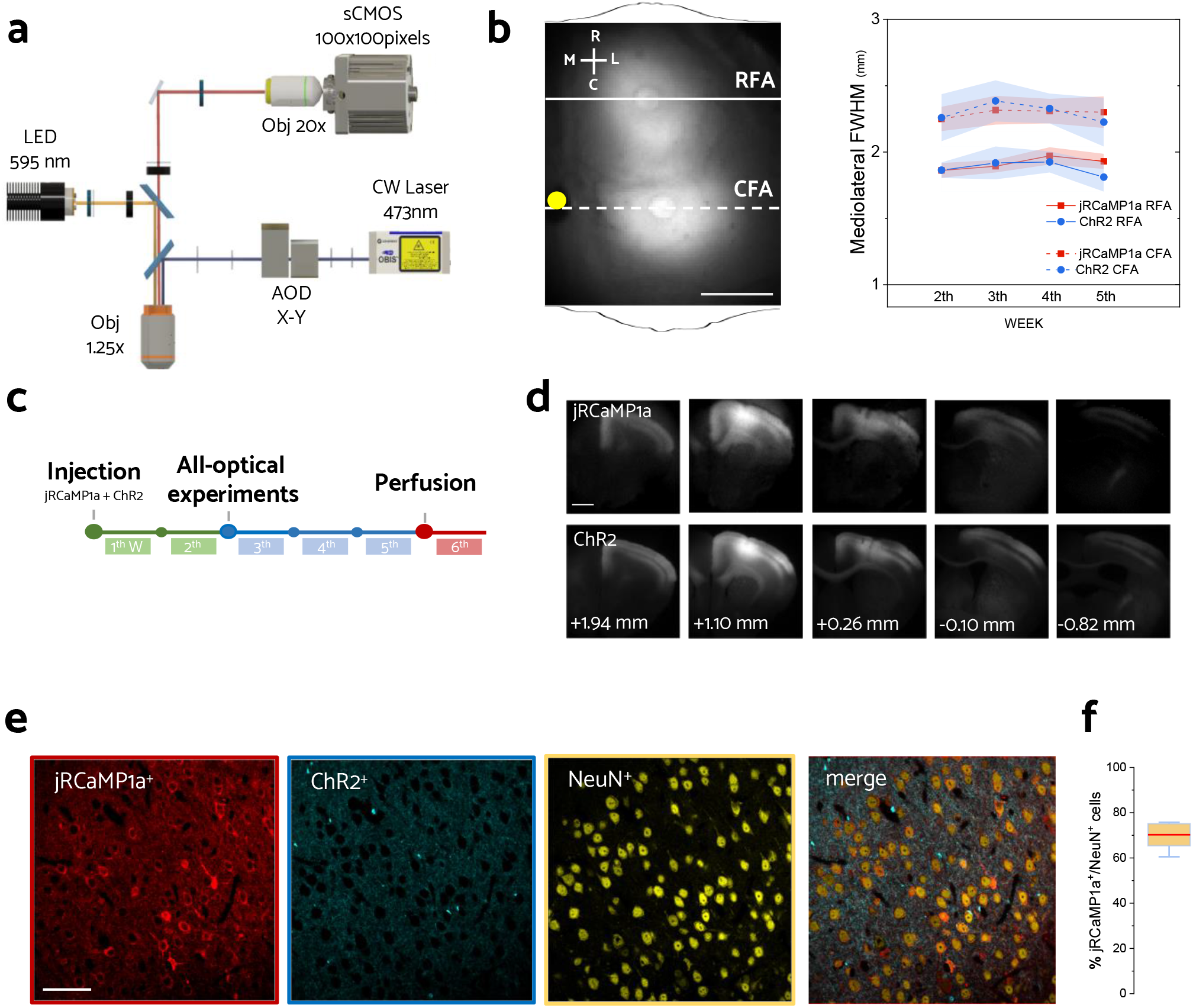
In vivo experimental design to perform parallel functional imaging and light-based motor mapping in awake mice. **(a)** Schematic representation of the double path wide-field fluorescence microscope. **(b)** *In vivo* long-term quantification of jRCaMP1a (red) and ChR2 (blue) spatial distribution along the mediolateral plane passing through RFA (n = 7; solid line) and CFA (n = 7; dashed line) injection sites. Yellow dot indicates bregma. Scale bar = 1 mm. White cross represents the rostro-caudal (R-C) and medio-lateral (M-L) axis **(c)** Experimental timeline. **(d)** Ex-vivo coronal slices showing the rostro-caudal transfection extension of jRCaMP1a and ChR2. Scale bar = 1 mm. **(e)** Representative immunohistochemistry images showing the neuronal expression of jRCaMP1a (red), ChR2 (blue) and NeuN (yellow), in the motor cortex. Scale bar 50 μm. **(f)** Quantification of the colocalization ratio jRCaMP1a+/NeuN+ (70,2 ± 4,9 %, n = 7). Error bars represent SEM.

### Wide-field imaging of jRCaMP1a does not induce ChR2 cross-activation

The all-optical approach we chose to visualize cortical activation during optogenetically-evoked complex movements combines blue-activated opsins and red-shifted GECIs^21–25^. To date, there are two main red-shifted GECI families, comprising the RGECO and RCaMP variants that are based on mApple and mRuby protein respectively. Despite RGECOs show higher Ca^2+^ affinity and larger dynamic range compared to RCaMPs, they exhibit significant photoactivation when stimulated with blue-light, thus hindering their combination with blue and green-activated opsins^21^. Indeed, most all-optical systems exploiting single-photon excitation critically suffer for crosstalk between imaging and photostimulation^19,21,26,27^. To assess the possible cross-activation of ChR2 during wide-field imaging of jRCaMP1a, we performed local field potential (LFP) recordings during an alternating on/off pattern of the imaging illumination path (supplementary fig. 1a). Illumination-triggered average of the LFP showed no significant differences in the normalized power content of the standard neurophysiological spectral bands (supplementary fig. 1b). This result suggests that there were not relevant alterations of the neuronal activity during imaging. Therefore, we evaluated whether the laser wavelength used for optogenetic stimulation affected the jRCaMP1a readout. Single laser pulses at increasing intensity were delivered in mice expressing (jRCaMP1a^+^/ChR2^+^) or lacking the optogenetic actuator (jRCaMP1a^+^/ChR2^-^) (supplementary fig. 1c). The results showed a clear asymptotic increase of the jRCaMP1a response in jRCaMP1a^+^/ChR2^+^ mice. Conversely, laser pulses in jRCaMP1a^+^/ChR2^-^ mice did not induce jRCaMP1a responses up to 20 mW (supplementary fig. 1c). the results demonstrate that our all-optical configuration wards off the crosstalk between imaging and photostimulation.

### Wide-field Imaging during light-based motor mapping reveals the presence of an activation threshold for complex movement execution

To map forelimb multi-joint movements, we performed optogenetic stimulation of several sites in the motor cortex identifying the locomotion-like movement (TAP; supplementary video I) and the grasping-like movement (GRASP; supplementary video II), (fig. 1a). To establish the movement-specific light-based motor maps (LBMMs), we initially stimulated the previously reported stereotaxic references for the Rostral Forelimb Area (RFA; + 2 mm AP, +1.25 mm LM) and the Caudal Forelimb Area (CFA; + 0.25 mm AP, + 1.5 mm LM), in order to induce GRASP and the TAP movements respectively (fig. 2a)^4,9,28^. Stimulus trains at increasing laser power were used to identify the threshold required to elicit a clear motor behavior within-subjects (fig. 2c-d). The laser pawer thresholds were then employed to design the GRASP and TAP LBMMs (fig. 2b). As previously reported, we found that GRASP and TAP LBMM stereotaxic references were centered in the RFA and CFA respectively^4,14^. The large range of power thresholds (1.3 - 13.2 mW), could be ascribed to both biological and ChR2 expression variability between subjects. On the contrary, the evoked calcium transients amplitude, that reflects the neuronal ensemble activation, showed limited variability among several animals (GRASP ΔF/F_peak_ = 15.5 ± 1 %; TAP ΔF/F_peak_ = 12.7 ± 1 %, n = 11; fig. 2e). These observations were further confirmed by the lack of a significant relationship between the stimulus intensities and the amplitude of the evoked calcium transients (fig. 2f). These results suggest a critical activation threshold for triggering complex movement execution.

**Fig. 2.**
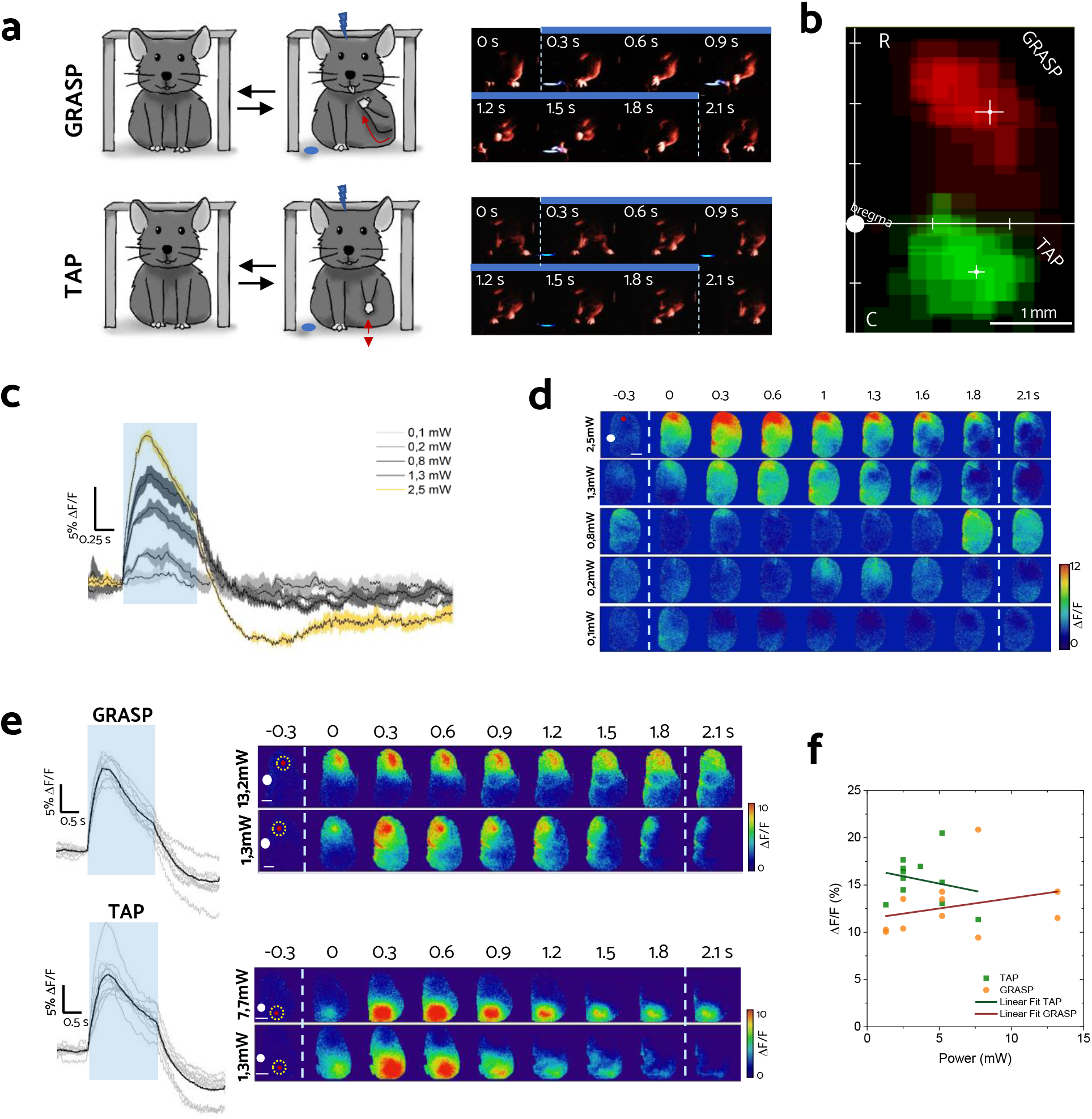
Wide-field Imaging during light-based motor mapping reveals an activation threshold for movement execution. **(a)** Left, representative cartoons describing the evoked movements. Blue dots are the reflected laser stimuli representation. Red arrows indicate movement trajectories. Right, example frames from behavior recording during grasping-like movement (top) and locomotion-like movement (bottom). **(b)** Average light-based motor maps for GRASP movement (red) and TAP movement (green). White crosses represent the maps centers of mass and cross-bar lengths represent SEM (GRASP RC = 1,8 ±0,2 mm; GRASP LM = 1,8 ± 0,2 mm; TAP RC = −1 ± 0,2 mm; TAP LM = 1,6 ± 0,2 mm; n = 8). **(c)** Representative average calcium responses to the optogenetic stimulus train (10 ms, 16 Hz, 2 s) at increasing laser powers. Yellow line represents the calcium response threshold associated with complex movement execution. Blue shadows represent the stimulation period. Shadows indicate SEM. **(d)** Representative wide-field image sequences of cortical activation at different laser powers. White dot indicates bregma. Red dot represents the site of stimulus. Dashed lines indicate the stimulus period. Scale bar = 1 mm. **(e)** Left panel. Calcium transients evoked at the minimum laser power (TAP, n=11; GRASP, n=11). Black line indicates average calcium transient. Right panel. Representative image sequences of cortical activation at minimum evoking power in two extremes (lower and higher power. Red dot represents the site of stimulus. Dashed lines indicate the stimulus period. Yellow dashed dots indicate the ROI where the calcium transients were calculated. White dot indicates bregma. Scale bar = 1mm. **(f)** Linear regression between power thresholds and evoked calcium transient amplitudes (TAP_intercept_ = 16.7 ± 1.8; TAP_slope_ = −0.3 ± 0.4; GRASP_intercept_ = 11.4 ± 1.7; GRASP_slope_ = 0.2 ± 0.2; n = 11).

### Movement-specific cortical functional connectivity is bounded to discrete modules

We wanted to clarify the intrinsic connections and reciprocal functional role of the motor cortices. To this aim, we analyzed the LBMM and the related cortical activation map for each movement category. First, we evaluated the calcium transients evoked in RFA, CFA and nomovement-evoking sites (supplementary fig. 2). The results showed no significant differences between RFA and CFA calcium transient amplitudes (supplementary fig. 2a), while both these areas showed a significantly higher peak amplitude compared with the surrounding nomovement-evoking sites (supplementary fig. 2b). This result suggests a stronger network activation in the movement-evoking areas compared to sites where no macroscopic movements were stimulated. To study the spatial distribution of the activated areas, we performed Maximum Intensity Projections (MIPs) of the cortical imaging stacks recorded during light-based motor mapping (fig. 3a). We defined the movement-specific activation map (MSAM) as the difference between the MIPs obtained within the LBMM boundaries and those obtained close outside those areas (supplementary fig. 2). Subsequently, to investigate the connectivity of the engaged cortical regions we analyzed the LBMM and MSAM spatial profiles (fig. 3a). The results showed that the MSAM area dimension and center of mass were comparable to the related LBMM (fig. 3b-c). Moreover, we evaluated the matching of the MSAM with the relative LBMM as the overlap between the maps of the two movement classes. The results showed that RFA and CFA activation networks (MASMs) were clustered in modules overlapping the respective movement cortical topography (LBMMs). Interestingly, RFA and CFA presented limited common activated areas as demonstrated by the low overlap of their MSAMs (fig. 3d and e). These results showed that during movement stimulation there was a large cortical activation confined to the relative movement representation area.

**Fig. 3.**
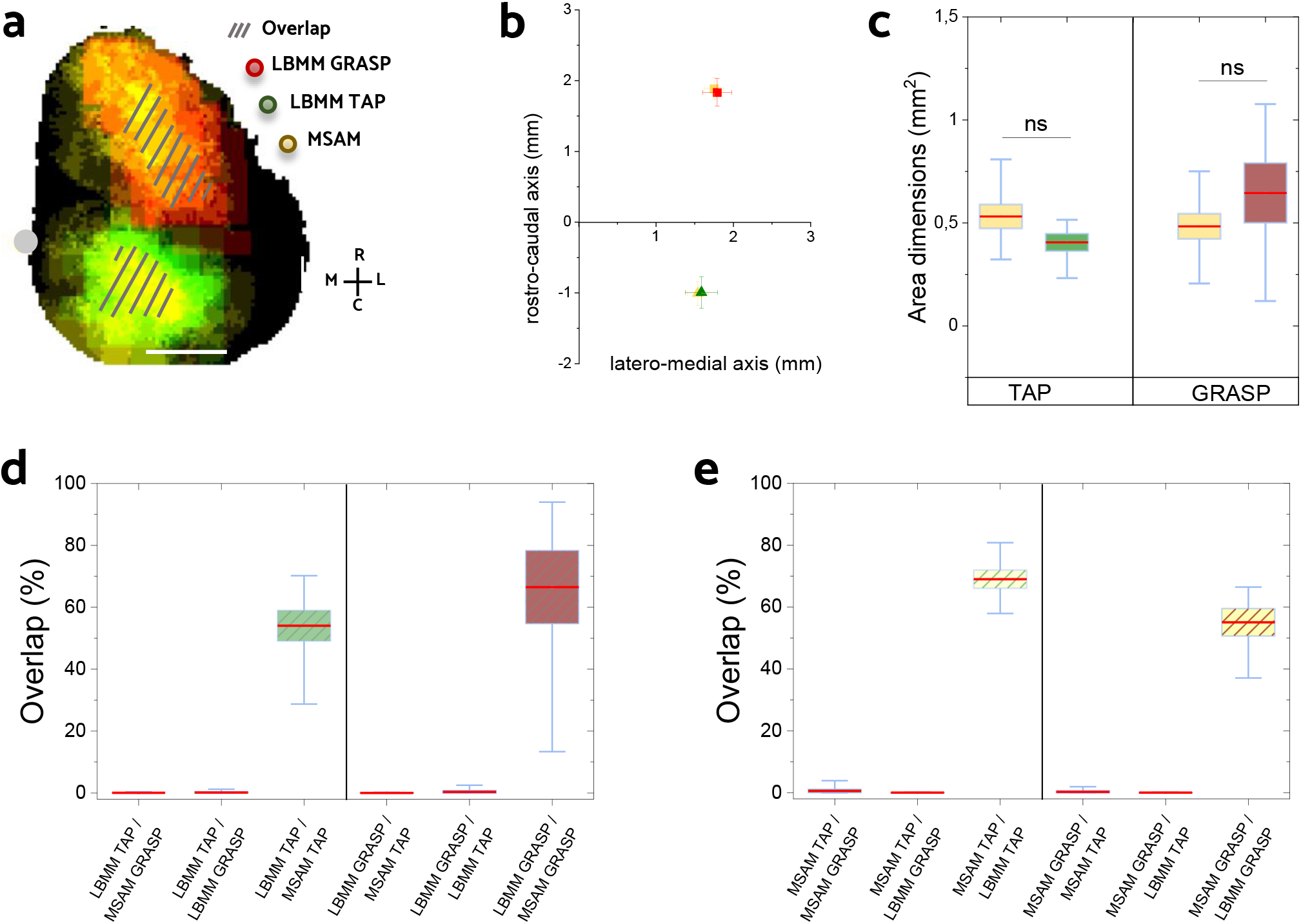
Movement-specific cortical functional connectivity is bounded to discrete modules. **(a)** Representative scheme of cortical movement representations (LBMM: TAP n = 8, red; GRASP n = 7, green) and the related average movement-specific activation maps (MSAM yellow: TAP n = 8; GRASP n = 7). Gray dot indicates bregma. Scale bar = 1 mm **(b)** Centers of mass of the LBMM (TAP MSAM: LM = 1.5 ± 0.1 mm; RC = −1.0 ± 0.2 mm; TAP LBMM LM = 1.6 ± 0.2 mm; RC = −1.0 ± 0.2 mm; n_TAP_ = 8. GRASP MSAM LM = 1.7 ± 0.1 mm; RC = 1.9 ± 0.2 mm; GRASP LBMM LM = 1.7 ± 0.2 mm; RC = 1.8 ± 0.2 mm; n_GRASP_ = 7). Colors as in (a). Cross-bar lengths represent SEM. **(c)** Quantification of the area dimensions of the LBMMs and the relative MSAM(TAP_MSAM_= 0.53 ± 0.06 mm^2^; TAP_LBMM_ = 0.40 ± 0.04 mm^2^;n_TAP_ = 8; GRASP_MASM_ = 0.48 ± 0.06 mm^2^; GRASP_LBMM_ =0.64 ±0.14 mm^2^; n_GRASP_ =7, two sample t-test). Comparison of the LBMMs **(d)** and the MSAMs **(e)** overlap (n = 7; see table 1).

### Identification of the Lateral Forelimb Area (LFA) as a distinct grasping representation module

Although optogenetically-evoked grasping-like and locomotion-like movements were elicited in areas corresponding to the RFA and CFA respectively, we observed that the GRASP LBMM was laterally and caudally stretched beyond the RFA (fig. 2b and 3a). In half of the examined subjects, this extended GRASP LBMM was separated into two distinct areas one frontal that matched the RFA and the other more caudal and closer to the lateral border of the CFA. We named this area the lateral forelimb area (LFA) (fig. 4a). Initially, we observed that LFA and RFA evoked calcium transients were similar (supplementary fig. 5), evidencing a comparable local neuronal response to the optogenetic stimulus. To understand whether the LFA represented a distinct module or it was an extension of the RFA network, we analyzed the LFA connectivity and its relation with the RFA and CFA. In animals presenting a single extended grasping representation area, we considered its caudal and lateral part as the GRASP LFA (see methods). Our results showed that, as for RFA and CFA, the LFA presented a clear matching of its LBMM and MSAM and their centers of mass were drastically separated from the others (fig. 4b and d). Remarkably, the limited overlap of the LFA MSAM with the maps of the other modules maps reinforced the hypothesis of the discrete functional area (fig. 4e-f). Interestingly, we observed a modest overlap between the MSAM over the LBMM of the LFA (fig. 4d), indicating wider connectivity that exceeded the LBMM borders of the LFA reaching distant areas (fig. 4a). These results show that the LFA is associated with specific cortical connectivity features different from that of RFA or CFA.

**Fig. 4.**
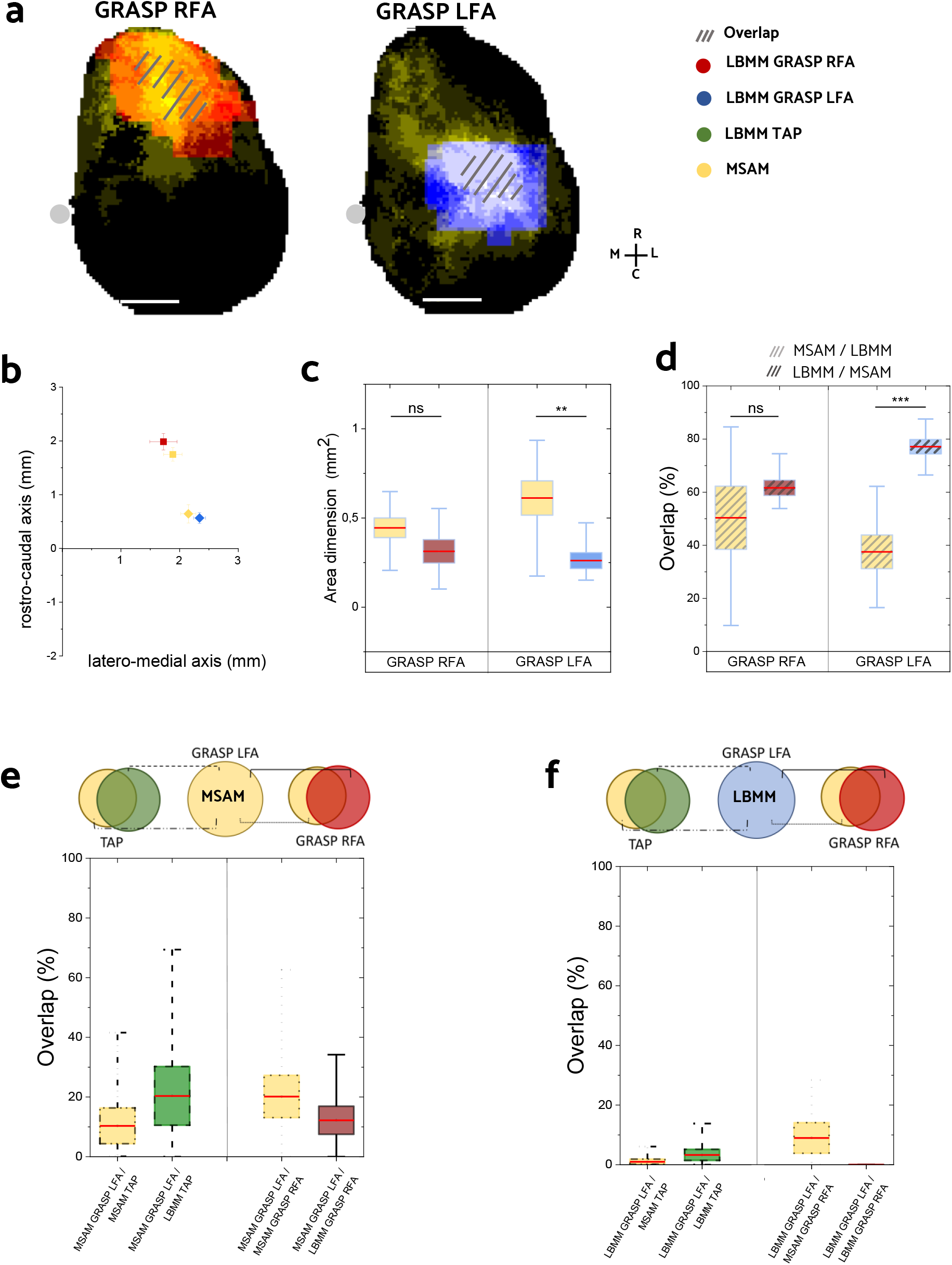
Identification of the Lateral Forelimb Area (LFA) as a distinct grasping representation module. **(a)** Representative schemes of the cortical movement representations (GRASP RFA in red; GRASP LFA in blue) and their related average MSAM (yellow) (n = 7). Gray dot indicates bregma. Scale bar = 1 mm. **(b)** Centers of mass of GRASP RFA LBMM (red), Grasp LFA LBMM (blue) and MSAMs (yellow) (LBMM RFA_RC_= 2.0 ± 0.2 mm; RFA_LM_= 1.7 ± 0.2 mm vs LFA_RC_= 0.6 ± 0.1 mm; LFA_LM_ = 2.3 ± 0.6 mm; MSAM RFA_RC_= 1.7 ± 0.1 mm; RFA_LM_= 1.9 ± 0.2 mm vs LFA_RC_ = 0.6 ± 0.2 mm; LFA_LM_ = 2.1 ± 0.1 mm; n = 7). **(c)** Quantification of the area dimensions of the LBMMs and the relative MSAM (GRASP RFA MSAM = 0.44 ± 0.06 mm2; GRASP RFA LBMM = 0.31 ± 0.07 mm2; GRASP LFA MSAM = 0.61 ± 0.09 mm2; GRASP LFA LBMM = 0.26 ± 0.04 mm2; n = 7, ** p<0,01 two sample t-test). **(d)** Quantification of the overlay between MSAMs and LBMMs per movement category (GRASP RFA_LBMM/MSAM_ =61 ± 3 %; GRASP LFA_LBMM / MSAM_ = 77 ± 2 %; GRASP RFA_MSAM / LBMM_ = 50 ± 12 %; GRASP LFA_MSAM / LBMM_ = 38 ± 6 %; n = 7, *** p<0,001 two sample t-test). Red lines indicates means, boxes show the standard error range, whiskers length represents the extreme data points. **(e)** Multiple comparison between GRASP LFA MSAM and the other movement category maps (n = 7; see table 2). **(f)** Multiple comparison between GRASP LFA LBMM and the other movement category maps (n = 7; see table 2).

### Grasping-like behaviors evoked in RFA and LFA exhibit similar kinematic profiles

At a glance, GRASP RFA and GRASP LFA movements presented similar profiles showing an initial forelimb displacement towards the midline followed by elevation to the mouth, which was often coupled with forepaw twisting and licking (fig. 2a; supplementary video II, III). Therefore, to examine in detail their trajectories we tracked the contralateral forelimb movements and performed kinematic analysis. Although GRASP LFA average trajectory was slightly wider compared to GRASP RFA (fig. 5c-d), the absolute maximum lateral displacement and elevation were similar (fig. 5e-f). Moreover, the movement onset time did not show a significant difference between all movement categories (fig. 5b). Conversely, TAP movements displayed completely different trajectories compared to GRASP. Indeed, the locomotion-like movement is rhythmic whereas the grasping-like is a discrete movement. TAP kinematics showed periodicity (fig. 5d), narrow medial-lateral displacement (fig. 5e) and weak elevation (fig. 5f). As expected, the stimulation of no-movement-evoking sites resulted in a remarkably reduced and non-specific forelimb displacement (fig. 5e-f). Overall, the similarity between GRASP RFA and GRASP LFA kinematics suggests that both cortical modules evoke the same movement.

**Fig. 5.**
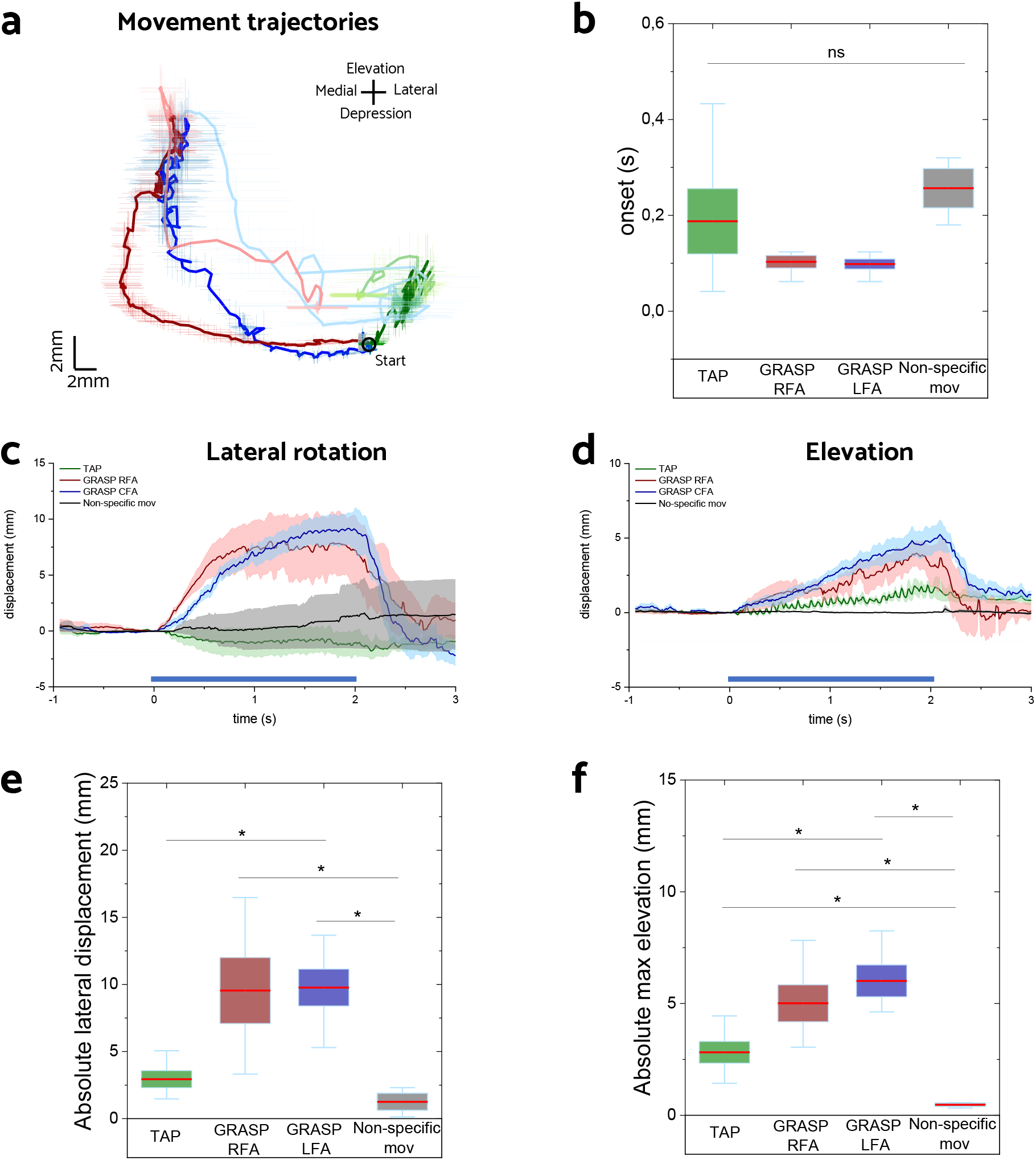
Grasping-like behaviors evoked in RFA and LFA exhibit similar kinematic profiles. **(a)** Reconstruction of the mean trajectories evoked by optogenetic stimulation of GRASP RFA (red), GRASP LFA (blue), TAP (green) and non-specific movement (light gray). Dark traces show the movement trajectory during the 2 s stimulus period. Light traces show the movement trajectory 1 s post stimulation. Black circle indicates the forelimb start point. Error bars represent SEM. **(b)** Box and whisker plots showing the onset time per movement type (TAP, 0.19 ± 0.07 s; GRASP RFA, 0.10 ± 0.01 s; GRASP LFA, 0.10 ± 0.01 s; n = 5; non-specific mov, 0.26 ± 0.04 s, n = 3; one-way ANOVA with post hoc Bonferroni test). **(c)** Medio-lateral forelimb displacement profiles. Dark traces represent the average medio-lateral displacement per movement category. Shadows indicate SEM. Blue line shows the stimulus period. **(d)** as in **(c)** mean elevation displacement along the y-axis. **(e)** Comparison of the absolute maximum displacement along the medio-lateral axis (TAP, 2.9 ± 0.6 mm; GRASP RFA, 9.5 ± 2.5 mm; GRASP LFA, 9.8 ± 1.4 mm; n = 5; non-specific mov, 1.2 ± 0.6, n = 3; P<0,05 one-way ANOVA with post hoc Bonferroni test). **(f)** Comparison of the absolute maximum elevation (TAP, 2.81 ± 0.48 mm; GRASP RFA, 5.01 ± 0.83 mm; GRASP LFA, 6.01 ± 0.71 mm; n = 5; non-specific mov, 0.47 ± 0.08, n = 3; p<0,05 one-way ANOVA with post hoc Bonferroni test).

### Cortical activity propagation analysis reveals movement-specific spatiotemporal patterns of activation

The spatial analysis highlighted that the optogenetically-evoked complex movements were associated with discrete modules of cortical activation. To investigate the spatiotemporal progression of this activation through cortical areas, we computed the cortical activity propagation map by ranking the time of activation of each pixel in the FOV (fig. 6, see also Methods). From a global (i.e., across the different animals) analysis of these maps, we obtained four polar plots (one for each stimulation condition) describing the spatial propagation direction. Results showed that during RFA stimulation there was a rapid activation of the area around the site of stimulus (green color, fifth rank or earlier) followed by a laterocaudal activation flow that largely preserves its spatial orientation, as shown in fig. 6a. A specular rostromedial flow of activation was observed during CFA stimulation (fig. 6b). Interestingly, LFA stimulation evoked a more complex pattern of cortical activation and the analysis pipeline hardly provides a clear direction for the activation flow (fig. 6c). In addition, no-movement-evoking stimulation sites showed a slower spread of activation compared with the complex movement-evoking sites, coupled with an unclear direction of propagation (fig. 6d). Overall, these results reveal that different complex movements are linked with specific spatiotemporal patterns of activity propagation. Interestingly, LFA showed peculiar propagation features, thus reinforcing the hypothesis that LFA relates to a specific grasping evoking module. In addition, the stimulation of movement-related cortical areas leads to a more marked flow of activation compared to no-movements-evoking areas, suggesting the persistence of more complex connectivity associated with movement execution.

**Fig. 6.**
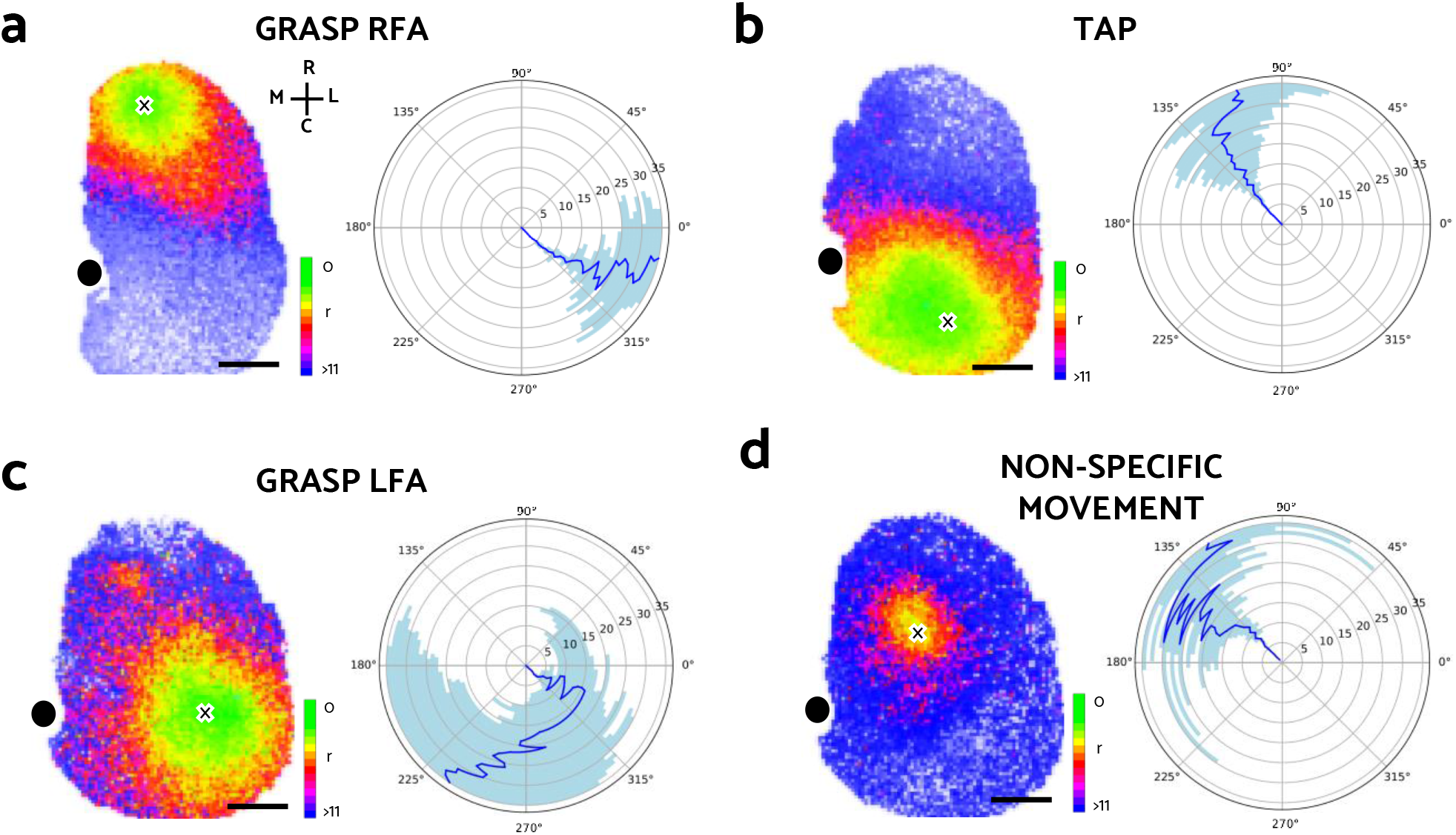
Cortical activity propagation analysis reveals movement-specific spatiotemporal patterns of activation. **(a)** Left panel. Average map of the spatiotemporal activity propagation during GRASP RFA stimulation. Right panel. Polar plot, centered on the stimulation site, showing the average propagation direction. Blue line represents the radius-dependent circular mean. Shadow represents the standard deviation. As in **(a)** spatiotemporal activity propagation maps and polar plots are shown in **(b)**, **(c)** and **(d)** for TAP, GRASP LFA and nonspecific movement stimulations respectively. Scale bars = 1 mm. Color bar = pixel ranks from 0 to <11.

### Excitatory synaptic block leads to disruption of connectivity features associated with complex movement interference

The spatiotemporal analysis suggested a progressive engagement of specific regions in the motor cortex. A common strategy to dissect the role of different functional nodes in a neuronal network is the pharmacological synaptic transmission block, in particular using glutamatergic transmission antagonists^14,29,30^. To investigate both the role of the local connectivity and the reciprocal role of each module during optogenetic-evoked movement execution, we performed a module-specific block of the excitatory synaptic transmission through topical application of the AMPA/kainate receptor antagonist 6-cyano-7-nitroquinoxaline-2,3-dione (CNQX) on the cortical surface^14,30,31^. The results showed that CNQX application in RFA reduces the extension of the GRASP RFA activation map (fig. 7b) while it does not significantly affect the calcium transient profiles (fig. 7c). Similar results were obtained by applying CNQX in CFA (fig. 7b-c). These results suggest an impairment of the local connectivity (a decrease in activation spreading) that does not affect the direct local response to the optogenetic stimulus. These results are in accordance with evidence showing that topical application of CNQX disrupts the cortical connectivity while preserving the direct activation of ChR2-expressing neurons^14^. Moreover, to analyze the effect of the module-specific block of the excitatory synaptic transmission on the activity propagation features, we compared the pixel rank distribution of an ROI overlapping the relative LBMM before and after the pharmacological block (fig. 7d-e). These results highlight an increase in both the median and the interquartile range (IQR) caused by the pharmacological connectivity interference suggesting a slower and more disorganized propagation of the cortical activity, respectively (fig. 7f). It should be noted that these results correlate with the behavioral outcome (fig. 7g). Indeed, as previously reported by Harrison et al., 2012, CNQX application leads to faults and distortions in complex movement execution, until the complete extinction of a recognizable complex movement^14^. Interestingly, CNQX application in RFA resulted in a block of GRASP execution while preserving the CFA-evoked TAP movement. The same result was obtained following the application of CNQX in CFA that resulted in a block of the TAP while maintaining a successful GRASP RFA expression. Taken together these results demonstrate that local excitatory synaptic inputs are required to control complex motor behaviors, and that GRASP RFA and TAP are controlled by two independent cortical modules.

**Fig. 7.**
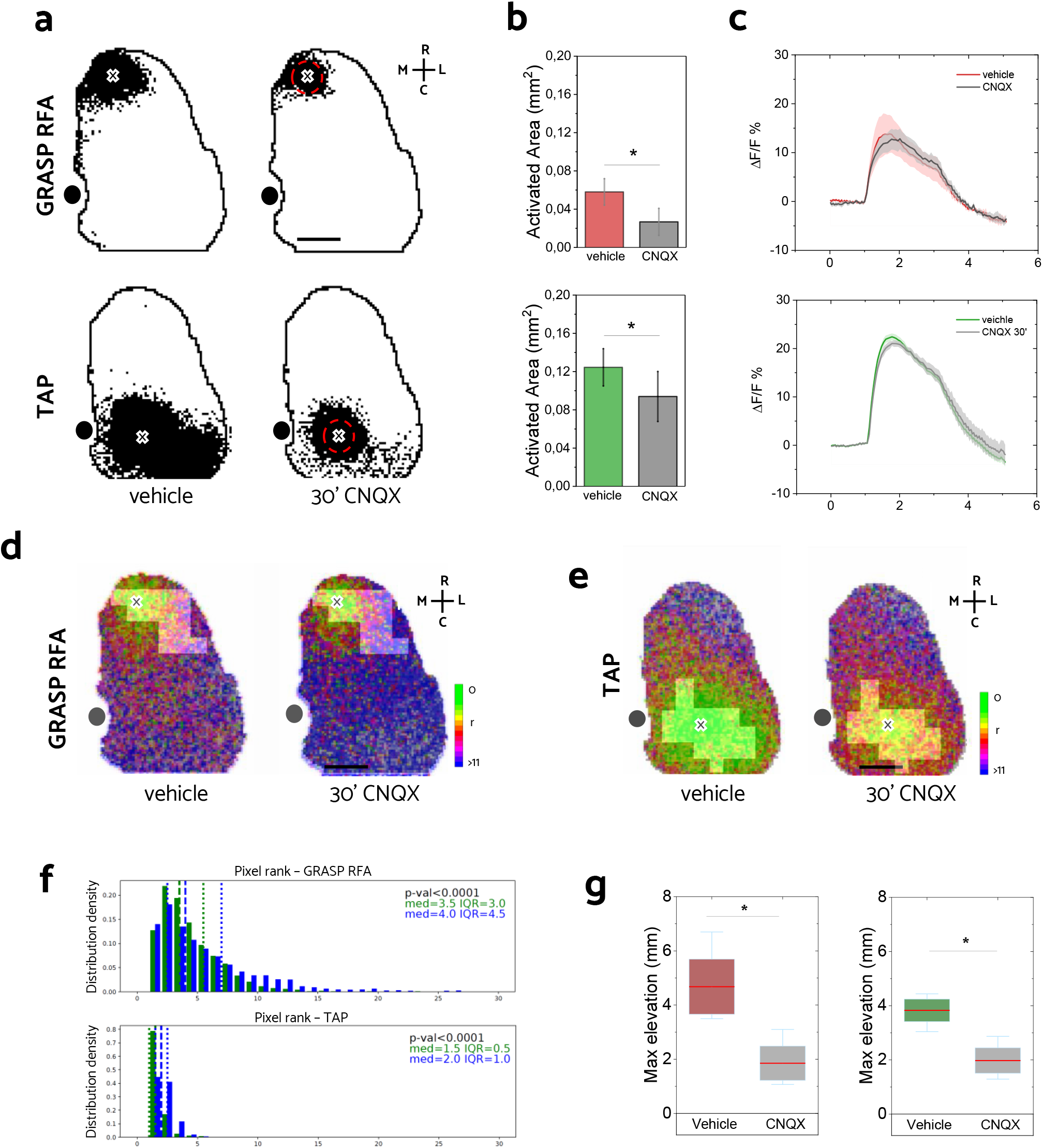
Excitatory synaptic block leads to disruption of connectivity features associated with complex movement interference. **(a)** Representative MIP showing optogenetically evoked cortical activation during vehicle (left) and CNQX topical application (right) in RFA (top) and CFA (bottom). Cross represents the stimulus site. Red dashed lines indicates the CNQX topical application site. **(b)** Quantification of the effect of CNQX topical application on MSAM extension in RFA (top) and CFA (bottom) (GRASP: vehicle = 0.05 ± 0.01 mm2 vs CNQX = 0.02 ± 0.01 mm2; n = 3, * p<0,05, paired sample t-test; TAP: vehicle = 0.12 ± 0.01 mm2 vs CNQX = 0.093 ± 0.026 mm2; n = 3, * p<0,05, paired sample t-test) **(c)** Averaged evoked calcium transient profiles in vehicle and following CNQX topical application in RFA (top) and CFA (bottom) (GRASP: vehicle 14.72 ± 4.20 ΔF/F vs CNQX 11.62 ± 3.66 ΔF/F; n = 3, paired sample t-test; TAP: vehicle 25.80 ± 1.31 ΔF/F vs CNQX 21.26 ± 0.63 ΔF/F; n = 3, paired sample t-test). Shadows represent SEM. Representative activity propagation maps of GRASP RFA **(d)** and TAP **(e)** showing the effect of CNQX topical application. Bright areas represent the LBMM. Scale bar = 1 mm. Color bar = pixel ranks from 0 to <11. **(f)** Pixel rank distribution of the region corresponding to the LBMM (bright) for GRASP RFA (top) and TAP (bottom), before (green) and after (blue) CNQX topical application (n = 3). Med = median, IQR = interquartile range, Wilcoxon signed-rank test. **(g)** CNQX topical application effect on movement kinematics. Comparison of the absolute left-forelimb maximum elevation in vehicle (color) and CNQX topical application (gray) in RFA (left) and CFA (right) (GRASP: vehicle 4.6 ± 1 mm vs CNQX 1.8 ± 0.6 mm; n = 3, * p<0,05, paired sample t-test; TAP: vehicle 3,8 ± 0,4 mm vs CNQX 1.9 ± 0.4 mm; n = 3, * p<0,05, paired sample t-test). Red lines indicates means, boxes show the standard error range, whiskers length represents the extreme data points.

### LFA-evoked grasping does not require RFA activation

Connectivity analysis suggested that the three identified functional modules activate distinct cortical regions during movement execution and the excitatory synaptic block experiments confirmed that GRASP RFA and TAP do not require mutual activation (fig. 7). In order to further study the relation between LFA and RFA, we stimulated the LFA during RFA pharmacological block (fig. 8). The results showed that there was no significant difference neither in the LFA MSAM dimension (fig. 8b) nor in evoked-calcium transients (fig. 8c). Moreover, we observed that the RFA pharmacological block did not modify the LFA spatiotemporal propagation features (fig. 8d-e). Interestingly, the preserved cortical connectivity of the LFA correlated with the successful execution of the GRASP LFA movement despite the block in RFA (fig. 8f). These results demonstrate that the LFA motor output does not require the RFA activation.

**Fig. 8.**
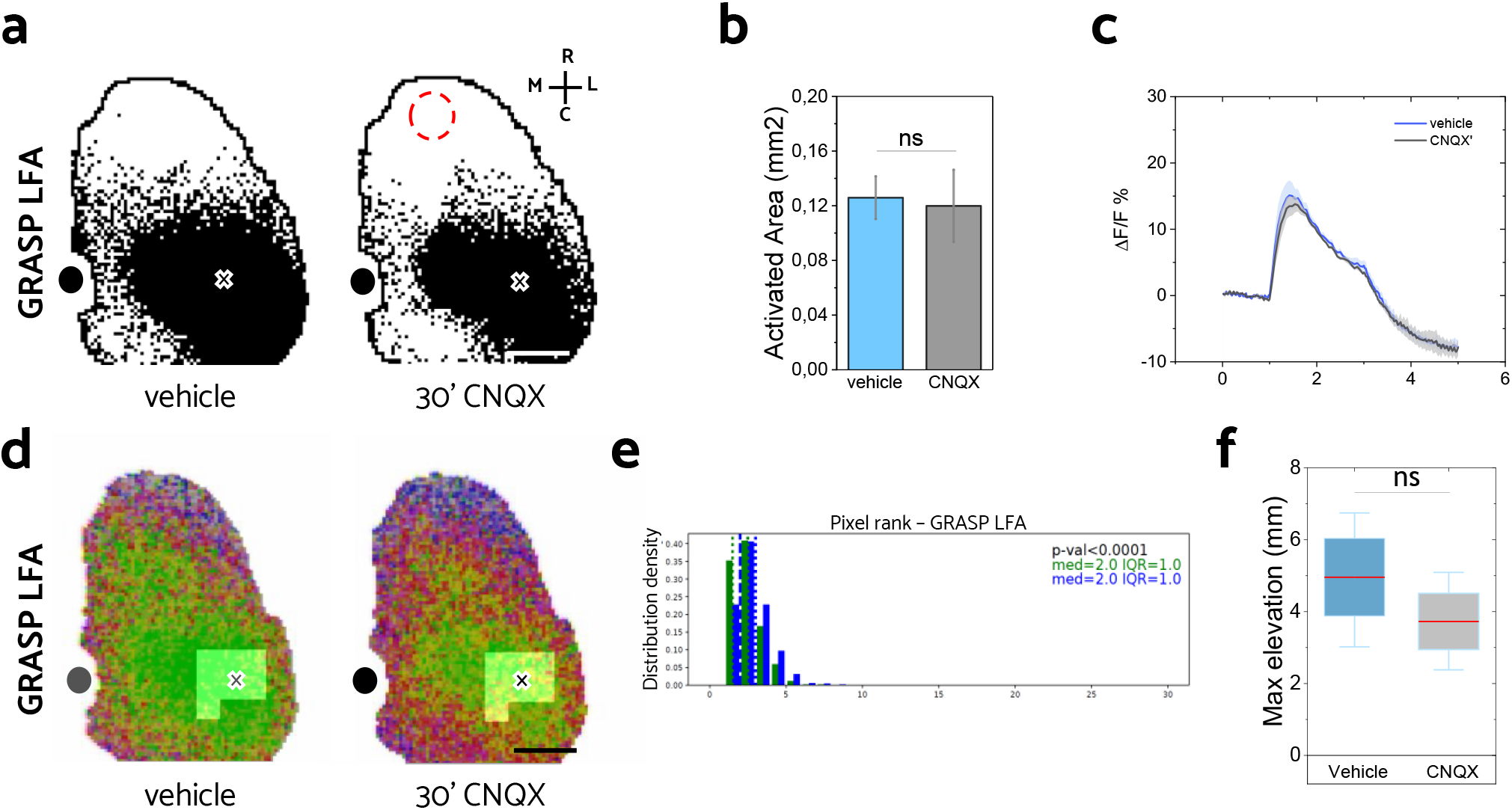
LFA-evoked grasping does not require RFA activation. **(a)** Representative MIPs showing optogenetically evoked cortical activation in LFA during RFA topical application of vehicle (left) and CNQX (right). Cross represents the stimulus site. Red dashed line indicate the CNQX topical application site. **(b)** Quantification of the effect of CNQX topical application in RFA on LFA MSAM extension (vehicle = 0.12 ± 0.01 mm2 vs CNQX = 0.11 ± 0.02 mm2; n = 3, paired sample t-test). **(c)** Averaged LFA evoked calcium transients profiles in vehicle and following CNQX topical application in RFA (vehicle 15 ± 2 ΔF/F vs CNQX 13 ± 1 ΔF/F; n = 3, paired sample t-test). Shadows represent SEM. **(d)** Representative activity propagation maps of GRASP LFA showing the effect of CNQX topical application in RFA. **(e)** Pixel rank distribution of the region corresponding to the LBMM (bright) for GRASP LFA, before (vehicle) and after CNQX topical application (n = 3). Med = median, IQR = interquartile range, Wilcoxon signed-rank test. **(f)** Effect of RFA CNQX topical application on movement kinematics. Comparison of the absolute left-forelimb maximum elevation following LFA stimulation in vehicle and RFA CNQX topical application (vehicle 4.9 ± 1,8 mm vs CNQX 3.7 ± 1.4 mm; n = 3, paired sample t-test). Red lines indicate means, boxes show the standard error range, whiskers length represents the extreme data points.

## DISCUSSION

Exploiting a large-scale all-optical method, we mapped the RFA and CFA activity during stimulated complex movement execution. Thanks to our novel approach we provided evidence for a segregated organization of RFA and CFA networks. The areas engaged during RFA and CFA stimulation were clustered over their optogenetic maps and did not overlap. Moreover, the spatiotemporal activity propagation analysis reveals movement-specific patterns. The pharmacological disruption of RFA or CFA inhibited the execution of the related movement while preserving the behavioral output and the activation features of the other areas, demonstrating their partitioned functional structure. Remarkably, we identified a second grasping representation area, functionally independent from the RFA and expressing distinct activation features, that we named Lateral Forelimb Area (LFA).

To causally investigate neuronal circuits, optogenetics has been paired with single- and two-photon fluorescence imaging in all-optical neurophysiology approaches^19,32^. There are still important limitations matching optogenetics and fluorescence imaging due to the large crosstalk between functional indicators and optogenetic actuators, caused by their spectral overlap. This phenomenon leads to spurious modulation of neuronal activity during imaging or stimulation artifacts in the readout channel, making it hard to develop a cross-talk free all-optical method^19,33,34^. Recently, Forli et al. took advantage of two-photon absorption for both jRCaMP1a excitation and ChR2 activation to manipulate and record cortical neuronal activity in anesthetized mice demonstrating this solution as the best option to drastically reduce crosstalk^23^. Here we extended this configuration to transcranial single-photon excitation in awake head-fixed mice. We initially evaluated the cross-activation of jRCaMP1a and ChR2 using visible light excitation. The electrophysiological analysis showed that the imaging excitation did not affect the LFP content, suggesting that there is no detectable neuronal activation during jRCaMP1a imaging (supplementary fig. 1a-b). This result is in line with previous evidence showing the absence of crosstalk using RCaMPs and ChR2^21–23^. In particular, using the patch-clamp technique, it has been demonstrated that one-photon wide-field illumination at 590 nm of cultured neurons expressing ChR2 prevents photocurrent generation, thus maintaining subthreshold potentials^23^. The second aspect we considered was the jRCaMP1a activation following blue-laser excitation. As shown in supplementary fig. 1c the laser stimulation did not change the jRCaMP1a fluorescence dynamics in mice lacking ChR2, demonstrating that optogenetic stimulation did not affect the neuronal activity readout. Besides, we optimized a transfection strategy to achieve a wide and stable expression of both the optogenetic actuator and the fluorescence reporter over one hemisphere, exploiting a double AAV viral vector injection. Consistently with our previous study^20^, we obtained a stable expression for both jRCaMP1a and ChR2 in the right hemisphere motor cortex (fig. 1). It should be noted that we did not observe bleaching of jRCaMP1a signal after prolonged exposure to our imaging sessions and after dozens of consecutive optogenetic stimulations (supplementary fig. 3; see methods). However, our method lacks optical sectioning, thus it is hard to control either the cortical layers targeted by the optogenetic stimulation or the exact source of the fluorescence signal given the spatial profile of the jRCaMP1a expression (fig. 1d). Therefore, further studies will be necessary to clarify these points.

Due to technical limitations, investigations of the cortical connectivity related to natural behaviors^17,18^ and motor mapping studies^4,35^ are largely confined to separated experiments. Previous research showed that a rich repertoire of complex movements can be evoked optogenetically-stimulating different sites in the mouse motor cortex^4,14^. In the present study, we developed a method to simultaneously analyze the cortical representation of complex movements and the related features of mesoscale activation. We focused on two forelimb movements, the discrete forepaw-to-mouth movement (GRASP) and the rhythmic locomotion-like movement (TAP) obtained by stimulating the RFA and CFA respectively^4,12,13^. According to the literature, the quantitative kinematic analysis revealed that the TAP trajectory exhibited rhythmic repetitions and a slight lateral displacement while the GRASP trajectory was mainly displaced in the mediolateral plane and exhibited a discrete elevation of the forelimb toward the mouth (fig. 5)^4,14^ Nevertheless, the observed movements were characterized by larger onset times compared to the literature, this discrepancy could be ascribed to both the lower stimulation frequency and the different animal models used.

In order to map the cortical topography of these movements, we evaluated the minimum laser power required to elicit a clear GRASP or TAP by gradually increasing the stimulus intensity. Results showed variable subject-specific power values. Interestingly, the related cortical activation exhibited lower between-subjects heterogeneity (fig. 2). This difference could be ascribed to the between-subjects variability in the opsin expression. Indeed, the efficiency of the optogenetic stimulation depends on the absolute protein expression whereas the imaging signal is normalized as relative fluorescence changes (ΔF/F), which reduces the impact of the expression variability on the readout. Once the subject-specific minimum laser power was identified, we designed the GRASP and TAP LBMMs. In accordance with previous studies, we found that GRASP and TAP LBMMs covered the RFA and CFA respectively (fig. 2b)^4,14^ Our novel all-optical tool allowed us to study the amplitude of the calcium transients evoked in GRASP and TAP LBMMs that were significantly different from those evoked in nomovement-evoking areas (Supplementary fig. 3b). This result suggests stronger intrinsic connectivity of the complex movement representation areas. Moreover, our spatial analysis revealed that the GRASP representation extended laterally beyond the RFA towards the forelimb somatosensory (FLS1) cortex (fig. 2b and 3a). Therefore, we characterized this lateralization identifying the LFA (fig. 4). Previously Bonazzi *et al*. mapped the motor cortex topography in anesthetized rats through ICMS, describing a lateral area expressing a hold-like forelimb movement, defined as the paw supination (the wrist and forearm turning toward the midline or the face)^10^. In the lateral part of the rat motor map, the authors showed that the hold-like movement is coupled with elevation and abduction movements that resemble the feature that we considered for grasping movement. Interestingly, also Harrison *et al*. exploiting the light-based motor mapping technique, observed a lateral extension of the forelimb abduction movement centered in the RFA. Moreover, previous studies reported that corticospinal motor neurons (CSN) can be found in RFA, CFA, and in a small circumscribed cluster in the secondary somatosensory cortex named PL-CFA^36,37^. At the spinal cord level, CSN axons from PL-CFA mainly overlap with the RFA-CSNs premotor neurons^36,38^, suggesting the control of the same group of muscles. Therefore, the LFA that we functionally characterized in this paper could refer to the group of neurons anatomically described in these previous works. Optogenetic stimulation of RFA and LFA lead to the generation of similar grasping behavior, therefore, we investigated the functional role of the LFA compared to the RFA, starting with the kinematic analysis of the respective movements. The results showed that GRASP RFA and GRASP LFA expressed comparable trajectories achieving equal elevation and displacement in the medial-lateral plane and displaying the same onset time (fig. 5). This result confirms that LFA and RFA exhibit the same motor output.

To study the connectivity hallmarks of GRASP RFA, GRASP LFA and TAP, we calculated the movement-specific activation maps (MSAMs). We found that the MSAMs were packed in segregated modules that largely overlapped the associated movements representation topography (fig. 3 and 4). Indeed, GRASP RFA topography and MSAM circumvent the TAP area and the relative MSAM. The same results were observed for the TAP movement. Remarkably, also the LFA activation map avoided the RFA- and CFA-relative LBMMs and MSAMs, displaying specific connectivity features (fig. 4). ICMS stimulation has been recently coupled with intrinsic signal optical imaging to reconstruct activation maps related to forelimb stimulated movements in squirrel monkeys^39^. The authors demonstrated that the intrinsic motor cortex connectivity matched the forelimb somatotopic representation in the primary motor cortex^39^. Our results are in line with evidence suggesting a segregated functional organization of CFA and RFA^4,11^, despite their mutual connections that may have a role in coordinating sequences of complex movements such as the reach to grasp behavior^40,41^. Interestingly, the spatial clustering of functionally correlated units in the motor cortex seems to be expressed across scales from neurons^42^ to entire functional areas^4^.

Spatiotemporal activity propagation features are pivotal aspects of the computation and communication between subsystems of the brain^43^. In the motor cortex, behaviorally relevant propagating patterns of cortical activation have been demonstrated to be necessary for movement initiation^44,45^. Therefore, we explored the spatiotemporal spreading of the neuronal activity during stimulated motor performances and we found movement-specific propagation patterns for the three motor regions (fig. 6). Our analysis revealed movement-specific orientation of the activity propagation, showing opposite directions for RFA and CFA. Conversely, LFA patterns exhibited more complex features rather than the fairly linear propagation observed for the other modules. These results reinforce the idea that LFA could represent a distinct GRASP representation.

To test the correlation between the simulated movements and the activity features observed, we performed module-specific inhibition of the excitatory synaptic transmission. This pharmacological tool allows an effective direct optogenetic stimulation of the targeted area while blocking its input connections, thus probing the role of the module-specific network in generating the forelimb movement. It has been demonstrated that topical application of the AMPA/kainate receptor antagonist CNQX on the cortical surface disrupts the optogenetic-evoked complex movement execution while preserving the direct activation of ChR2-expressing neurons^14^. Accordingly, our results show that the RFA pharmacological inactivation interferes with the GRASP execution while retaining the ability to evoke the TAP movement and the specular phenomenon was observed during CFA inhibition (fig. 7), supporting the idea of two functionally independent modules^4,14^. Moreover, we reported that CNQX application leads to a significant reduction of the MSAM extension associated with slower and more disorganized patterns of local propagation (fig. 7), highlighting that the activation features we described reflect the module-specific network activity linked to movement execution. These results sharpen the idea that direct activation of corticospinal projections is not sufficient to stimulate full movement performance, thus confirming the pivotal role of the cortical synaptic inputs. Further studies will be necessary to understand the contribution of the recurrent cortico-cortical circuits^46,47^ or subcortical loops^48^ in complex movement control and to define whether the modules found in the motor cortex and their related inputs can be organized as central pattern generator networks, in which the activation of a group of neurons can be sufficient to elicit an entire motor engram^49,50^. Moreover, we observed that during RFA inactivation the LFA behavioral output and all its activation features were preserved (fig. 8), demonstrating that the GRASP LFA expression is not affected by the RFA network, thus suggesting that the two grasping modules are parallelly organized.

To the best of our knowledge, this is the first application of a large-scale all-optical method in awake mice. The experimental paradigm we developed represents a powerful approach to causally dissect the cortical connectivity, reaching its full potential in experimental settings where it is not possible to record behavioral outputs, for instance in the study of non-motor cortical regions or the investigation of different brain states and pathologies with altered level of consciousness i.e., sleep, anesthesia or coma^51^. Exploiting this method, we raised evidence for a segregated functional organization of CFA and RFA and we identified a new forelimb representation area. Further studies will be necessary to define the ethological role of the LFA and its engagement in voluntary movements.

## METHODS

### Virus injection and intact-skull window

All experiments were performed in accordance with the guidelines of the Italian Minister of Health (aut. n. 871/2018). C57BL/6J adult mice (6-12 months) of both sexes were anesthetized with isoflurane (3% for induction, 1-2% for maintenance) and placed in a stereotaxic apparatus (KOPF, model 1900). Ophthalmic gel (Lacrilube) was applied to prevent eye drying, body temperature was maintained at 36°C using a heating pad and lidocaine 2% was used as local anesthetic. The skin and the periosteum were cleaned and removed. Bregma was signed with a black fine-tip pen. To achieve widespread expression of both jRCaMP1a and ChR2 over the right hemisphere, small holes were drilled at two coordinates (AP +2.0 mm, ML +1.7 mm; AP −0.5 mm, LM +1,7 mm from bregma). A 500 nl volume of mixed viruses (pGP-AAV9-syn-NES-jRCaMP1a-WPRE.211.1488 and pAAV9-CamKII-hChR2(H134R)-Cerulean, 1×10^13^ GC ml^-1^, CliniSciences, 250 nl respectively) was pressure-injected through a pulled glass micropipette at one depth per site (−0.5 mm ventral from dura surface) using an electrically gated pressure injector (Picospritzer III–Science Products™, n 3 Hz, ON 4 ms) for a total volume of 1 μl per mouse. A custom-made aluminum head-bar placed behind lambda and a cover glass implanted on the exposed skull were fixed using transparent dental cement (Super Bond C&B – Sun Medical). After the surgery, mice were recovered in a temperature- and humidity-controlled room, with food and water ad libitum for two weeks before recordings.

### Wide-field microscopy setup

Wide-field imaging and optogenetic stimulation were performed using a custom-made microscope with two excitation sources to simultaneously excite the opsin (ChR2-cerulean) and the calcium indicator (jRCaMP1a)^52^. The excitation source for jRCaMP1a was a red-light beam of emitting diodes (595nm LED light, M595L3 Thorlabs, New Jersey, United State) and the excitation band was selected by a bandpass filter (578/21 nm, Semrock, Rochester, New York, USA). The light beam was deflected by a dichroic mirror (606nm, Semrock, Rochester, New York, USA) to the objective (2.5x EC Plan Neofluoar, NA 0.085) towards the skull. The excitation source for single-photon stimulation of ChR2 was a continuous wavelength (CW) laser (λ = 473 nm, OBIS 473 nm LX 75mW, Coherent, Santa Clara, CA, USA). The excitation beam was overlaid on the imaging pathway using a second dichroic beam splitter (FF484-Fdi01-25 × 36, Semrock, Rochester, New York, NY, USA) before the objective. The system has a random-access scanning head with two orthogonally-mounted acousto-optical deflectors (DTSXY400, AA Opto-Electronic, Orsay France). The jRCaMP1a fluorescence signal emitted was collected through a band-pass filter (630/69, Semrock, Rochester, New York, USA) and focused by a tube lens (500 nm) on the sensor of a demagnified (20X objective, LD Plan Neofluar, 20×/0.4 M27, Carl Zeiss Microscopy, Oberkochen, Germany) high speed complementary metal-oxide semiconductor (CMOS) camera (Orca Flash 4.0 Hamamatsu Photonics, NJ, USA). The camera acquired images at a resolution of 100 by 100 pixels covering a quadratic field-of-view of 5.2 by 5.2 mm^2^ of the cortex.

### Wide-field imaging in awake mice

14 days after the injection, head-fixed imaging sessions were performed for three consecutive weeks. An animal-specific field of view (FOV) template was used to manually adjust the imaging field daily. Each imaging session consisted of 5-10 s of recording in resting-state followed by the stimulus train (2 s) and 30 s of imaging after the stimulus (sampling rate: 50 Hz). The waiting time for consecutive sessions was 3 minutes per animal. LED light intensity was 4 mW after the objective.

### Transcranial optogenetic stimulation

Laser stimulation patterns were generated using two orthogonally-mounted acousto-optical deflectors controlled by a custom-written LabView 2013 software (National Instruments). A reference image of the FOV was used to target the laser beam on a selected cortex area.

### Single-pulse laser stimulation

consisted of one pulse (10ms ON) repeated 8 times in one imaging session at different laser power (0.22 - 1.3 - 2.5 - 5.2 - 7.7 - 13.2 mW, after the objective).

### The stimulus train

consisted of 2 s, 16 Hz, 10ms ON. For laser power calibration experiments the laser power used were: 1,3 - 2,5 - 5,2 - 7,7 - 13,2 mW. For light-based motor mapping, connectivity studies and pharmacological inhibition laser power was the minimum power required to evoke movements (from 1.3 mW to 13.2 mW).

### Light-based motor map (LBMM)

The LBMMs for locomotion-like (TAP) and grasping-like (GRASP) movements were obtained in separate experiments. A virtual grid (14 x 14, 364 μm spacing) was superimposed on the animal-specific FOV template using Fiji^53^. A stimulus train was then delivered in a random order single time for all sites of the grid. The left forepaw position during imaging sessions was monitored using a camera equipped with a red illumination light focused on the forepaw and not interfering with imaging. Forelimb movements were evaluated by two different expert observers and visually categorized as (i) grasping-like movements: contralateral forepaw was closed, the wrist turned and moved toward the mouth (ii) locomotion-like movements: contralateral forelimb was retracted and lifted at least twice, simulating a walking movement (iii) no-movements and movement interference: the absence of at least one movement criterion during the stimulus. The average LBMM was created by aligning three points in the FOV (bregma and injection sites) in 7 animals per movement category.

### Optogenetic of the Lateral Forelimb Area (LFA) LBMM

4 out of 8 animals presented a discrete LFA and RFA light-based motor map. The average of these LFA LBMM, with a 100% threshold (total overlap for high restriction), was used as a mask for identifying the LFA border in those animals that presented a unified light-based motor map.

### Video tracking analysis

A machine vision camera (PointGrey flir Chamaleon3, CM3-U3-13Y3C-CS) was orthogonally set 100 mm in front of the mouse to evaluate the left forelimb movement induced by contralateral optogenetic stimulation (frame rate 100 Hz). The camera acquired images at a resolution of 800 by 600 pixels covering a field-of-view of 24 by 18 mm^2^ of the cortex (0.03 mm per pixel). Visible illumination light at 630 nm was focused on the left forepaw, to avoid imaging interference. Five individual trains per movement category for each animal were filmed (n_mice_ = 5; n_trains_ = 5). Videos were analyzed using the ImageJ plugin AnimalTracker, obtaining XY coordinates of the forelimb in each frame from a starting point (Ref.^54^ for details). To compare evoked complex movements, we analyzed the tracked forelimb mediolateral displacement, the elevation and the speed (mm/s) for each train.

### In-vivo local field potential recording

Local field potentials (LFPs) were recorded in the center of the transfected area. Glass pipettes were used to avoid light-induced artifacts during the electrophysiological recordings and were filled with a 2 M NaCl solution. The electrode was advanced, through a little hole in the skull, into the motor cortex L5 (800 μm from the dura surface) using a motorized micromanipulator (EXFO Burleigh PCS6000 Motorized Manipulator). Signals were amplified with a 3000 AC/DC differential amplifier, sampled at 10 kHz, highpass filtered at 0.1 Hz and lowpass filtered at 3 kHz. A reference and ground screws were placed on the occipital bone. LFP signal was recorded during a randomly activated pattern of led ON / led OFF (2 seconds each, 4.5 mW). As a control, an optogenetic single-pulse stimulus was delivered close to the pipette tip, resulting in a fast downward deflection, indicating that ChR2 was effectively transfected and functioning.

### Preprocessing of imaging data

Images were analyzed with ImageJ and OriginPro (OriginLab 2017). Frames displaying artifactual excitation of skull autofluorescence were removed (2 out of 3 frames) and interpolated. A daily individual mask was created using the maximum intensity projection of the first imaging session (baseline). Masks were thresholded twice the mean value of the non-transfected hemisphere. For each imaging session, the fluorescence ratio change (ΔF/F_0_) was calculated averaging the first 50 frames before the stimulus onset (baseline fluorescence signal; F_0_).

### Calcium data analysis. In vivo quantification of jRCaMP1a and ChR2 spatial distribution

The full width at half-maximum (FHWM) of spatial fluorescence profile *in vivo* was evaluated during the third and fourth weeks after injection. FWHM was calculated on the average of the three brightest frames acquired in the resting state imaging session, over two parallel lines that crossed the injection sites in the mediolateral plane.

### Single-pulse correlation

was performed during the third and fourth weeks after injection. Single-pulse laser stimulations were delivered to the cortex region with the maximum level of ChR2 and jRCaMP1a expression. The consequently evoked calcium response dynamics (time series) were extracted from a region of interest (ROI, area 0,24 mm^2^) placed over the stimulation site.

### Power calibration

The optogenetic stimulus was delivered in the center of the light-based motor map. The stimulus train was repeated 3 times at increasing laser power to select the minimum power required to evoke GRASP and TAP movements. Calcium dynamics (time series) were extracted from a ROI (area 0,24 mm^2^) placed over the site of stimulation.

### Movement specific calcium map

An imaging stack was recorded for each site of the grid stimulated during the light-based motor mapping. For each acquisition (14 x 14, 364 μm spacing), the Maximum-Intensity Projection (MIP) was obtained and subsequently grouped by evoked-movement category in (i) GRASP RFA movement, (ii) GRASP LFA movement, (iii) TAP movement. Since light-based motor maps for TAP and GRASP were obtained in separate experiments, there were two different groups of no-movement: (iv) GRASP-related no-movement and (v) TAP-related no-movement. An average maximum activity value based on all MIPs was then calculated for GRASP RFA/LFA and TAP categories, and half of that value has been used for thresholding the MIPs of all groups. Thresholded MIPs were averaged and an additional threshold of 2x standard deviation (SD) was applied, obtaining the average activation map for all five experimental groups. Finally, in order to obtain movement-specific activation maps (MSAMs), the non-specific movement average activation maps were spatially subtracted from those related to GRASP RFA, GRASP LFA and TAP. The spatial overlap between the MSAM and the related LBMM was then assessed and quantified as a percentage of the total dimension of both the maps involved.

### Spatiotemporal propagation analysis

Spatiotemporal propagation analysis was performed with custom-made Python (Python Software Foundation, Beaverton, Oregon, U.S.A.) scripts. In a pre-processing step, image sequences were spatially masked and frames containing the laser stimulations were manually eliminated and replaced with their temporal linear interpolation. Then a Gaussian smoothing was performed along the temporal dimension (with a standard deviation for Gaussian kernel equal to one) before computing the ΔF/F_0_ signal. F_0_ was set as the average fluorescence value observed before the first laser stimulus. Pixels were identified as active if the maximum value of the ΔF/F_0_ signal after the first laser stimulus was larger than both the average value and the double of the standard deviation value, computed in both cases before the first stimulus. In active pixels, the time frame corresponding to the first crossing of a pixel-based threshold was used to identify the timing of the response to the laser stimulus. The threshold was set as twice the standard deviation value of the ΔF/F_0_ signal computed before the first laser stimulus. For imaging acquisitions pertaining to CNQX manipulation, the standard deviation values used to identify the active pixels and to define the timing thresholds were computed solely on the data acquired before CNQX administration (vehicle). For the data acquired after CNQX administration, the average of the standard deviations computed before CNQX administration was employed. In all cases, the timing values were then rank-transformed. The rank values of the active pixels related to the same animal and the same condition were averaged and the standard deviation was computed, while the non-active pixel values were discarded. Starting from these averaged results, for data related to CNQX manipulation, rank distributions were computed in a region of interest (ROI) overlapping the LBMMs. The distribution medians before and after CNQX administration were then compared using Wilcoxon signed-rank test. Moreover, to summarize the distribution characteristics, their interquartile ranges were computed alongside the medians. Finally, to trace the propagation direction, for each averaged result, pixels placed along a circumference centered on the laser-stimulated area and with varying radius were selected, discarding nonactive or masked pixels. For each circumference radius, the averaged rank distribution was computed and values composing its first quintile were sub-selected. Then the circular mean^55^ of the angular position (relative to the circumference center) of these values was computed. Finally, all the computed circular means were used to calculate the final, radius-dependent, circular mean and circular standard deviation^55^.

### Pharmacology

For pharmacological interference experiments, a small craniotomy (1 mm diameter) was performed on the region of interest. For the animals that underwent pharmacological inhibition of both GRASP and TAP, the experiments were performed in a 3day separate section. The craniotomies were then sealed with Kwik-seal (World Precision Instrument) after the experimental sections. Glutamate receptor antagonist CNQX 1 mM (C127 Sigma-Aldrich) and vehicle (physiological solution containing 0.01 % DMSO) were applied to the craniotomy and the solutions were replenished (at the same concentration) every 10’ to compensate for tissue drying. Stimulation sessions were performed every 10’.

### Immunohistochemistry

Four weeks after injection, mice were perfused with 20–30 ml of 0.1 M PBS (pH 7.6) and 150 ml of 4% paraformaldehyde (PFA). Brain coronal slices (100 μm thick) were cut with a vibrating-blade microtome (Vibratome Series 1500–Tissue Sectioning System). Slices were washed with PBS and incubated in PBS/0.3% Triton X-100 containing 1% bovine serum albumin (BSA) for 60 min while shaking at room temperature (RT). Then, slices were washed with PBS/0.1% Triton X-100 (T-PBS) and incubated with the primary antibody NeuN (1:200, Sigma, ABN78) in T-PBS for 1 day at 4°C while shaking. Then, slices were washed with T-PBS and incubated with anti-rabbit fluorescent Alexa 514 antibody (1:250, ThermoFisher, A-31558) in T-PBS for 2 h at RT while shaking. Finally, slices were washed and mounted on a glass slide. Imaging was performed with a confocal laser scanning microscope (CLSM, Nikon Eclipse TE300, with the Nikon C2 scanning head), equipped with a Nikon Plan EPO 60× objective, N.A. 1.4, oil immersion). The setup was equipped with 408 nm, 488 nm and 561 nm lasers to simultaneously excite ChR2, Alexa 514 and jRCaMP1a, respectively. A triple-band dichroic mirror 408/488/543 was used for simultaneous 3-channel fluorescence imaging. Emission filters were 472/10 nm, 520/35 nm and 630/69 nm.

### Statistics

All statistical analysis was performed in OriginLab 2018 except for the spatiotemporal propagation analysis that was performed with custom-made Python (Python Software Foundation, Beaverton, Oregon, U.S.A.) scripts. Data are shown as mean ± s.e.m. Parametric tests were used only after verifying for normality of the data employing Shapiro-Wilk and Kolmogorov-Smirnov tests. The error bars and shadows in graphs represent the s.e.m. In the box charts, the red line corresponds to the mean, the box shows the standard error range, whiskers lengths are the extreme data points. Student’s t-test was employed for every comparison concerning two samples and its paired version was used for paired data (fig. 7b and c; fig. 8b and c). For spectral band multiple comparisons in supplementary fig. 1b, two-way (variables: illumination status and bands) ANOVA was used. For multiple comparisons in Figure 6b,e,f (for onset, distance and elevation values, respectively) and in Supplementary figure 5a (for average calcium transients), one-way ANOVA was used and Bonferroni correction was applied for post-hoc t-tests. The level of significance was set at *p < 0.05, **p < 0.01, and ***p < 0.001.

## ACKNOWLEDGMENTS

This research has been supported by the European Union’s Horizon 2020 research and innovation Framework Programme under grant agreements N. 945539 (HBP-SGA3), N. 785907 (HBP-SGA2) and from the EU program H2020 EXCELLENT SCIENCE - European Research Council (ERC) under grant agreement n. 692943 (BrainBIT).

This research has also been supported by the Italian Ministry for Education, University, and Research in the framework of the Advance Lightsheet Microscopy Italian Mode of EuroBioimaging ERIC.

---

**Supplementary fig. 1.**
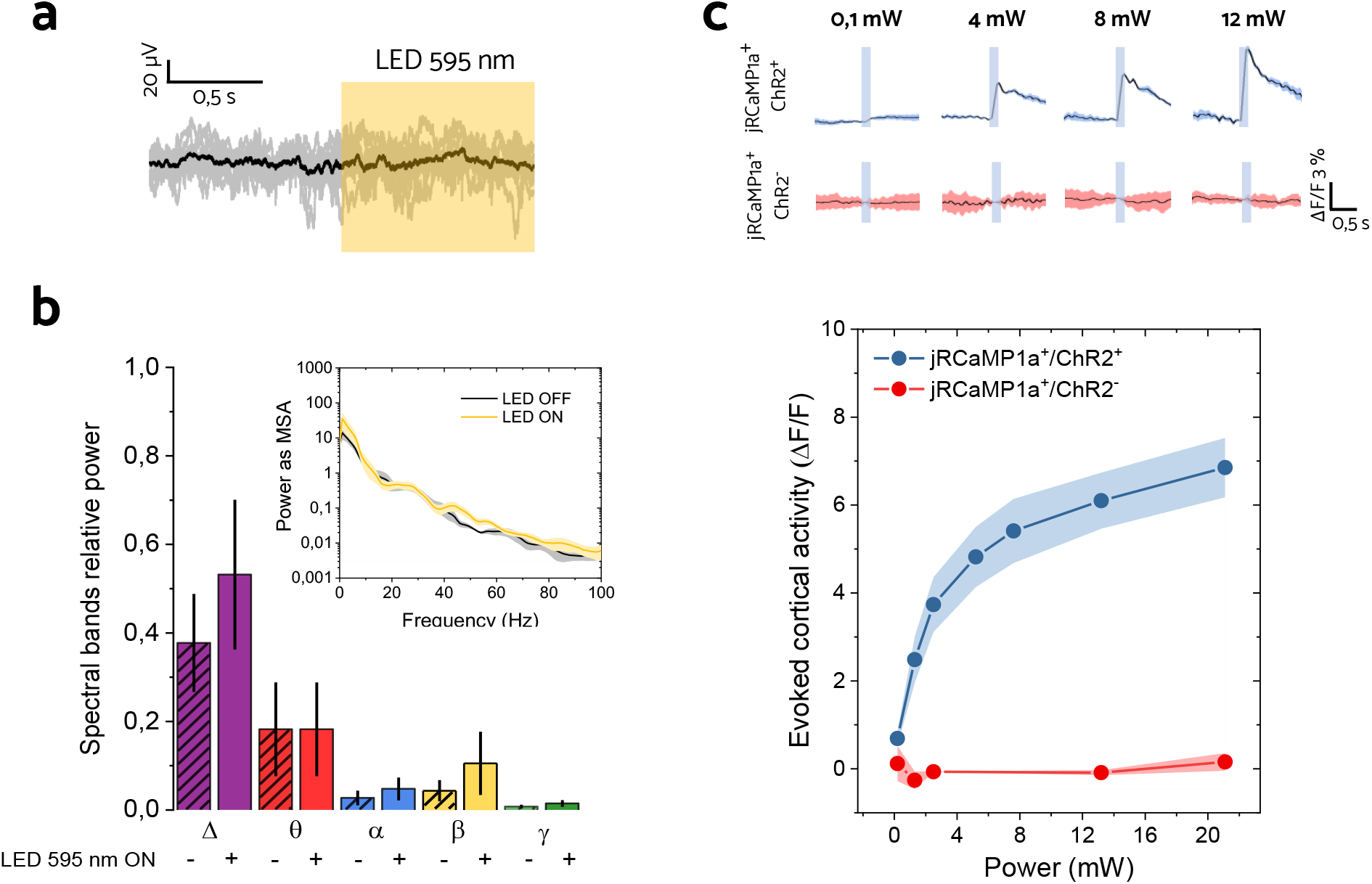
Wide-field imaging of jRCaMP1a does not induce ChR2 cross-activation. **(a)** Representative traces of the LFP signal (gray; n = 10) during 2 s of stimulus. Black line represents the mean signal. **(b)** Quantification of the spectral band relative power during 2 s of peri-stimulus period (n = 4 mice; 10 stimuli per mice). Columns represent the averaged relative power for LFP frequency bands (**Δ; θ; α; β; γ**) during the dark period (−2 - 0 s; patterned) and the LED illumination period (0 - +2 s; monochrome), n_mice_ = 4, n_stimuli_ = 10, two-way ANOVA with post-hoc Bonferroni test, Data are presented as mean ± SEM). Inset, representative averaged LFP power spectrum of 10 stimuli in dark period (grey) and LED illumination (Yellow). Shadows represents SEM. **(c)** Top panel. Representative traces showing the average calcium response (black) and the SEM (shadows) at increasing laser power (0,1 - 10 mW; 20 ms laser illumination) in mice expressing jRCaMP1a + ChR2 (blue) and only jRCaMP1a (red). Bottom panel. Correlation between evoked calcium activity and single pulse laser power in mice expressing jRCaMP1a+/ChR2+ (blue; n = 8) and jRCaMP1a+/ChR2- (red; n= 2). Shadows represents SEM.

**Supplementary fig. 2.**
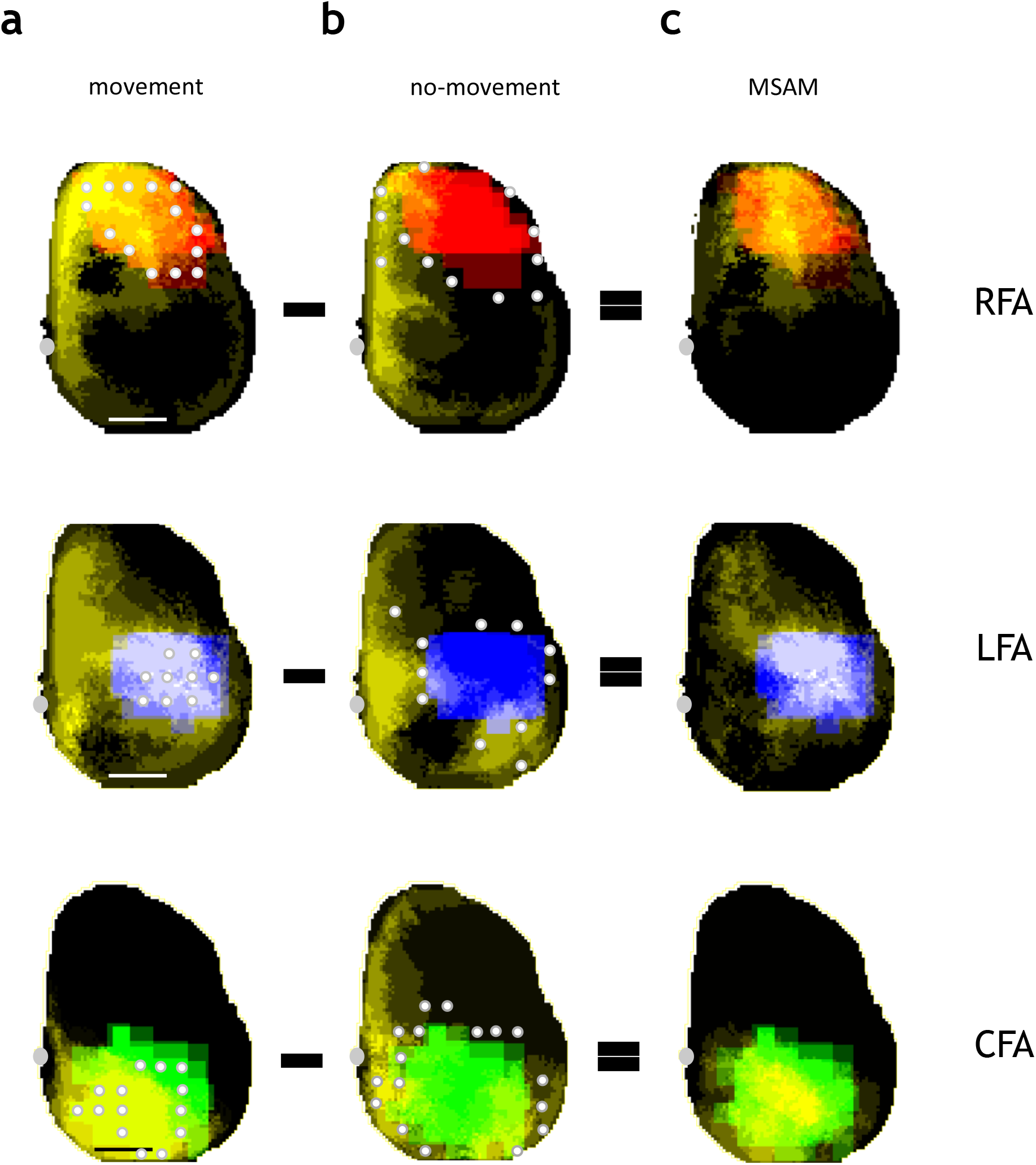
Movement-specific activation maps (MSAMs) processing. Average activation maps (yellow) obtained stimulating points (white dots) within the LBMM (colored areas) **(a)** and outside the LBMM **(b). (c)** Representative movement-specific activation maps (MSAMs) obtained by subtracting maps obtained in (b) to those obtained in (a).

**Supplementary fig. 3.**
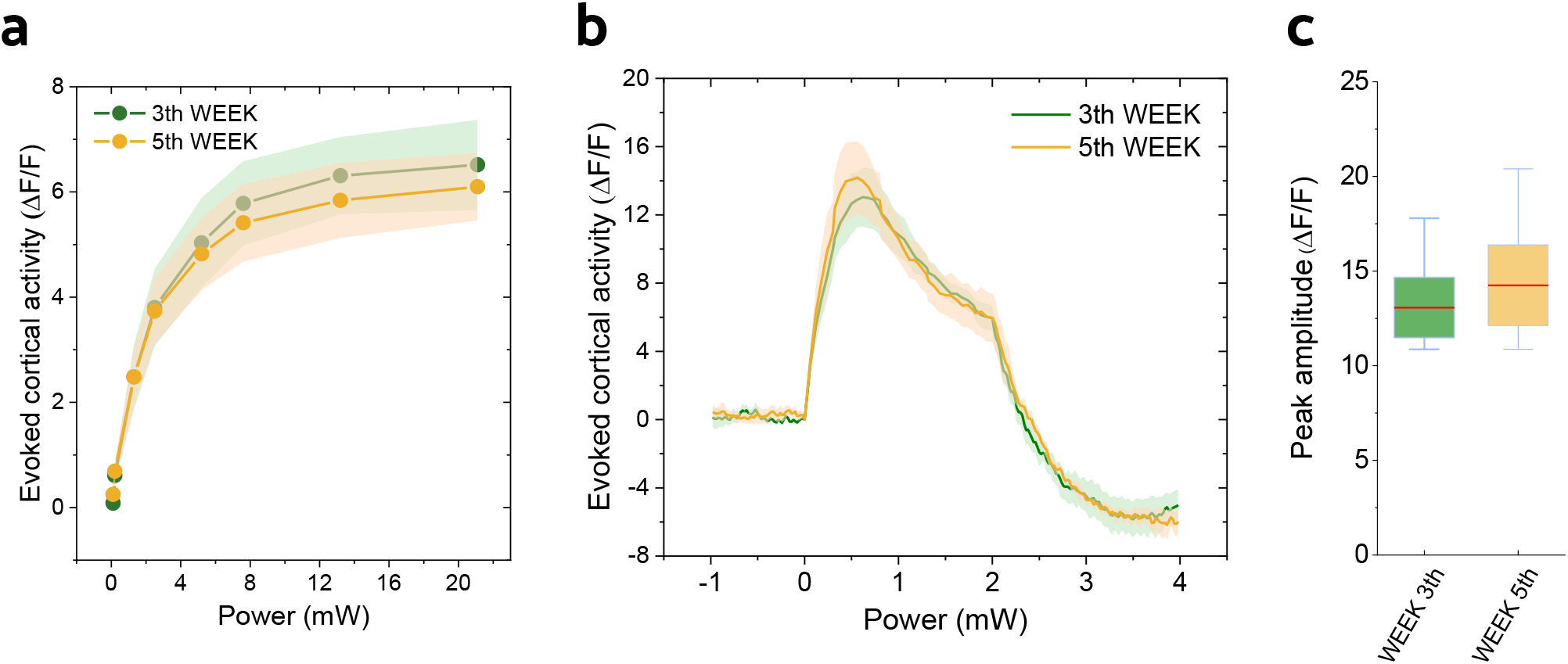
Optogenetically-evoked calcium transients show long-term stability. **(a)** Correlation between single laser pulse (10 ms) intensities and the calcium transient responses 3 and 5 weeks post-infection. **(b)** Calcium transients evoked by optogenetic stimulus trains (20 ms, 16 Hz; 2s) showing long-term profile stability. **(c)** Quantification of the stimulus train-evoked calcium transient amplitudes

**Supplementary fig. 4.**
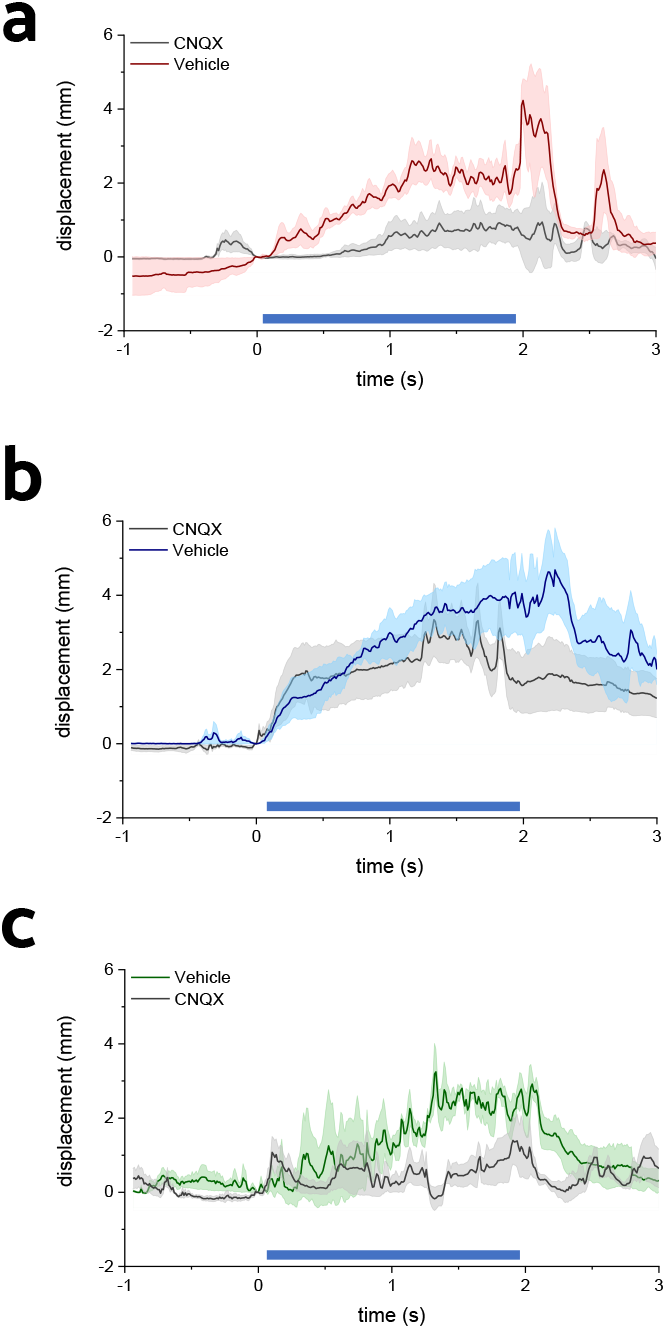
Excitatoty synaptic transmission block disrupts the complex forelimb movement execution. **(a)** Forelimb displacement during GRASP RFA before and after CNQX topical application (n = 3). Blue line indicates stimulation period. Dark traces represent the average and shadows represent SEM. **(b)** and **(c)** show the same analysis for GRASP LFA and TAP respectively.

**Supplementary fig. 5.**
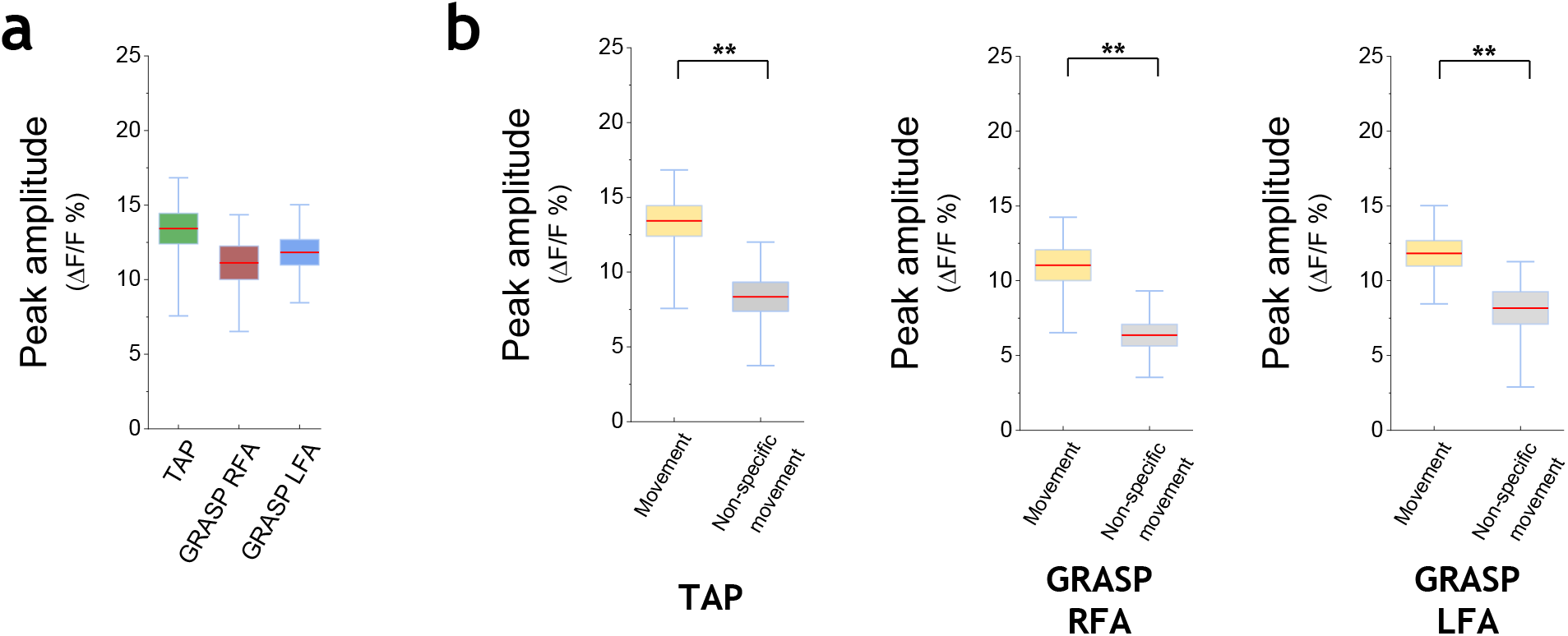
Comparison of average calcium transient properties between movement categories. **(a)** Comparison of evoked calcium transient amplitudes between movement categories (TAP 13.4 ± 1.0; GRASP RFA 11.1 ± 1.1 ΔF/F; GRASP LFA 11.8 ± 0.8 ΔF/F; n = 7, oneway ANOVA with post hoc Bonferroni test) **(b)** Comparison of the calcium transient amplitudes obtained stimulating within the LBMM (Movement) and outside the LBMM (non-specific movement) per movement classes (TAP: Movement = 13.4 ± 1.0 ΔF/F vs Non-specific movement 8.3 ± 0.9 ΔF/F; GRASP RFA: Movement = 11.1 ± 1.0 ΔF/F vs Non-specific movement 6.3 ± 0.7 ΔF/F; GRASP LFA: Movement = 11.8 ± 0.8 ΔF/F vs Non-specific movement 8.1 ± 1.0 ΔF/F; n = 7, ** p<0,01 two sample t-test).

